# Cold acclimation reprograms hepatic lipid composition toward n-3 HUFAs to uncouple adipose-derived lipid flux from steatosis

**DOI:** 10.64898/2026.03.27.714471

**Authors:** Victor Azevêdo Vieira, Gabriel Salerno Costa, Gabriela Segura Gonzales, Raphael Campos Guimarães, Cristiane dos Santos, Raul Gobato da Costa, Marcella Ramos Sant’Ana, Camila de Oliveira Ramos, Murilo Henrique Anzolini Cassiano, João Manoel Alves, Nayara Pereira, Paulo Henrique de Melo, Tiago Tomazini Gonçalves, Isabella Bonilha, Caio Fábio Baêta Lopes, Luiz Gustavo Gardinassi, Marcos Y. Yoshinaga, Luciane C. Alberici, Andrei C. Sposito, Dennys E. Cintra, Samir Softic, C. Ronald Kahn, Jing X. Kang, Tathiane M. Malta, Marcelo A. Mori, Yu-Hua Tseng, Luiz Osório Leiria

**Affiliations:** Department of Pharmacology; Ribeirao Preto Medical School, University of São Paulo, Ribeirão Preto, Brazil; Department of Cellular and Molecular Biology; Ribeirao Preto Medical School, University of São Paulo, Ribeirão Preto, Brazil; Center for Research in Inflammatory Diseases, CRID, University of São Paulo, São Paulo, Brazil; Immune Health Laboratory, “Regulation of host responses and immune health” (IRL2029), French National Centre for Scientific Research (CNRS) and Ribeirão Preto Medical School, University of São Paulo, Ribeirão Preto, Brazil; Section on Integrative Physiology and Metabolism, Joslin Diabetes Center, Harvard Medical School, Boston, USA; Department of Biochemistry and Tissue Biology, Biology Institute, State University of Campinas, Campinas, Brazil; Laboratory for Lipid Medicine and Technology, Department of Medicine, Massachusetts General Hospital and Harvard Medical School, Boston, USA; Omega-3 and Global Health Institute, Boston, USA; Department of Pediatrics and Gastroenterology, University of Kentucky College of Medicine, Lexington, USA; Laboratory of Nutritional Genomics, State University of Campinas, Limeira, Brazil; Department of Medicine, Faculty of Medical Science, State University of Campinas, Campinas, Brazil; Department of BioMolecular Sciences, School of Pharmaceutical Sciences of Ribeirão Preto, University of São Paulo, Ribeirão Preto, Brazil; Institute of Physical Activity and Sport Sciences, Universidade Cruzeiro do Sul, São Paulo, Brazil; PinguisLab, São Paulo, Brazil; Ribeirão Preto Nursing School, University of São Paulo, Ribeirão Preto, Brazil; Obesity and Comorbidities Research Center (OCRC), University of Campinas, Campinas, Brazil; Institute of Environmental, Chemical and Pharmaceutical Sciences - Federal University of São Paulo (UNIFESP). São Paulo, Brazil

**Author notes:** These authors contributed equally for the study (Victor Azevêdo Vieira, Gabriel Salerno Costa, Gabriela Segura Gonzales, Raphael Campos Guimarães).

## Abstract

While cold exposure drives lipid flux from adipose tissue to the liver, this enhanced inter-organ crosstalk does not result in sustained hepatic steatosis during prolonged acclimation, indicating that factors beyond lipid flux shape metabolic outcome. To interrogate this adaptation, we performed integrated lipidomic and metabolic profiling across tissues and circulating lipoproteins over the course of cold acclimation. We showed that cold acclimation induces systemic reprogramming of lipid quality in mice, characterized by enrichment of n-3 highly unsaturated fatty acids (HUFAs) as a consequence of upregulation of fatty acid desaturases (FADS1 and 2) in the liver and white adipose tissue, thus increasing hepatic n-3/n-6 ratio. Cold-induced increase in n-3 HUFAs cause the suppression of SCD1-mediated desaturation, thus yielding a depletion of monounsaturated fatty acids (MUFAs) in the liver, along with the suppression of lipogenic markers. Notably, this high-HUFA/low-MUFA lipid signature is present in both hepatic free fatty acid and triglyceride pools, indicating that lipid remodeling occurs upstream of triglyceride synthesis. Lipidomic analysis revealed that the remodeled triglycerides are incorporated into very-low-density and intermediate density lipoproteins (VLDL and IDL), thereby propagating hepatic lipid reprogramming to the circulation. Thus, by selectively increasing endogenous n-3 HUFA availability, cold adaptation suppresses hepatic DNL and MUFA-driven triglyceride assembly, buffering lipid accumulation despite sustained fatty acid influx and reshaping systemic lipid distribution with potential cardiometabolic impact.

## Introduction

Chronic exposure to cold activates a coordinated physiological program in mammals known as non-shivering thermogenesis, which increases energy expenditure, mobilizes energetic substrates, and enhances heat production primarily through the actions driven by thermogenic fat depots, namely brown adipose tissue (BAT) and inducible beige fat^1–4^. Beyond heat production, cold acclimation improves systemic glucose and lipid homeostasis^5,6^, while elevated BAT activity in humans is associated with reduced risk of cardiovascular disease and type 2 diabetes^7,8^. Lipids play a central role in this adaptation, functioning both as mitochondrial fuels and as bioactive signaling molecules^5,9–12^. Cold exposure robustly stimulates adipose tissue lipolysis, thereby promoting lipid flux from adipose tissue to peripheral organs, including the liver^13–18^. Notably, a similar adipose–liver lipid flux is observed in obesity and insulin resistance, where it contributes to hepatic steatosis and metabolic dysfunction^18^.

However, despite a pronounced increase in adipose-derived lipid flux during cold acclimation, the liver does not develop sustained steatosis. Instead, cold exposure induces a transient hepatic lipid accumulation that is resolved during adaptation^16,17^. These observations suggest that the metabolic consequences of adipose-derived lipid flux are context dependent and raise the possibility that lipid metabolism and/or partitioning, rather than flux alone, determines hepatic and systemic outcomes. Growing evidence supports the concept that lipid composition, including chain length, degree of unsaturation, and double-bond positioning and oxidation, shapes metabolic signaling pathways in the liver^11,19^. Fatty acids can be broadly classified as saturated (SFAs), monounsaturated (MUFAs), and polyunsaturated (PUFAs), including highly unsaturated fatty acids (HUFAs), based on their degree of unsaturation. SFAs are synthesized *de novo*, whereas MUFAs are generated via Scd1-mediated desaturation. In contrast, PUFAs are essential fatty acids that cannot be synthesized de novo in mammals and typically contain two to three double bonds, while HUFAs contain four or more double bonds. These fatty acids are derived from dietary precursors that can be further elongated and desaturated by ELOVL and FADS enzymes to generate long-chain PUFAs and HUFAs. Importantly, these enzymatic pathways operate in parallel and are dynamically regulated, enabling continuous remodeling of fatty acid composition^20^.

Previous study showed that young adult humans exposed acutely to cold exhibited increased levels of HUFAs in the form of circulating free fatty acids (FFA) and triglycerides (TGs) in a lipolysis-dependent manner^21^, suggesting that cold-induced adipose lipolysis may selectively shape the quality of lipids delivered systemically^21^. Nevertheless, whether cold acclimation engages coordinated inter-organ mechanisms that control fatty acid desaturation and trafficking to regulate the composition and accumulation of hepatic and circulating lipid pools remains unknown.

Here, we identify a coordinated lipid remodeling program induced by cold acclimation in mice. By integrating temporal fatty-acid profiling and molecular/metabolic assessment, along with untargeted lipidomics of hepatic and circulating lipoprotein fractions, we show that cold exposure selectively enriches n-3 HUFAs within both FFA and TG pools in the liver, concomitant with suppression of monounsaturated fatty acid production and reduced SCD1-dependent desaturation activity. These changes indicate that n-3 HUFAs are not merely trafficked through the liver, but are metabolically active in reshaping hepatic lipid synthesis and storage. As a consequence, the liver preserves HUFA-enriched triglycerides and selectively packages them into VLDL particles at the expense of monounsaturated fatty acids (MUFAs)-containing species. Collectively, these findings uncover a cold-responsive adipose–liver axis in which n-3 HUFAs act as regulators of hepatic lipid remodeling, thereby controlling the quality of circulating lipids during cold adaptation.

## Results

### Cold adaptation induces opposing lipid remodeling in adipose tissue and liver

First, we sought to characterize the overall lipid profile of inguinal and perigonadal white adipose tissue (ingWAT and pgWAT), BAT, and the liver of male C57BL/6J mice acclimated in thermoneutrality (30 °C) or cold (5 °C) for seven days (Fig. 1a). Experiments were done in fed condition. We chose to focus on total fatty acids aiming to capture the most abundant lipid species that either exert biological effects or serve as precursors for oxylipin biosynthesis. Cold exposure induced pronounced lipid remodeling across all analyzed tissues, consistent with the PCA separation between conditions (Fig. 1b). While adipose tissue depots clustered more closely together, the liver exhibited the greatest divergence relative to the other tissues (Fig. 1b). It caught our attention that the liver lipid profile was characterized by an overall increase in HUFAs and a concomitant decrease in MUFAs in cold condition (Fig. 1c-d). Interestingly, white adipose depots displayed divergent profile, as we found that cold elevated MUFAs content in these tissues and reduced HUFAs in pgWAT while in ingWAT it was not modified (Fig. 1d). These fluctuations in HUFA levels in WAT may reflect enhanced lipolysis under cold exposure, leading to continuous release of these fatty acids. Consistent with the reduction in MUFAs, we also observed an accumulation of saturated fatty acids (SFAs) in the liver (Fig. 1e), likely reflecting a lower enzymatic desaturation rate, as indicated by the reduced SCD1-dependent desaturation index (MUFA/SFA ratio) (Fig. 1g). PUFA levels were not changed in the liver, while they were reduced in white adipose depots (Fig. 1f). Importantly, we found an increase in the omega-3 (n-3) fatty acids content in the liver in response to cold acclimation, while no changes were found in n-6 content, resulting in higher n-3/n-6 ratio (Fig. 1h-j). This result contrasted with the overall reduction of n-3 content in adipose depots under cold (Fig. 1h-j). Noteworthy, higher n-3/n-6 is usually associated with better metabolic and anti-inflammatory profile in the liver^11^.

**Figure 1.**
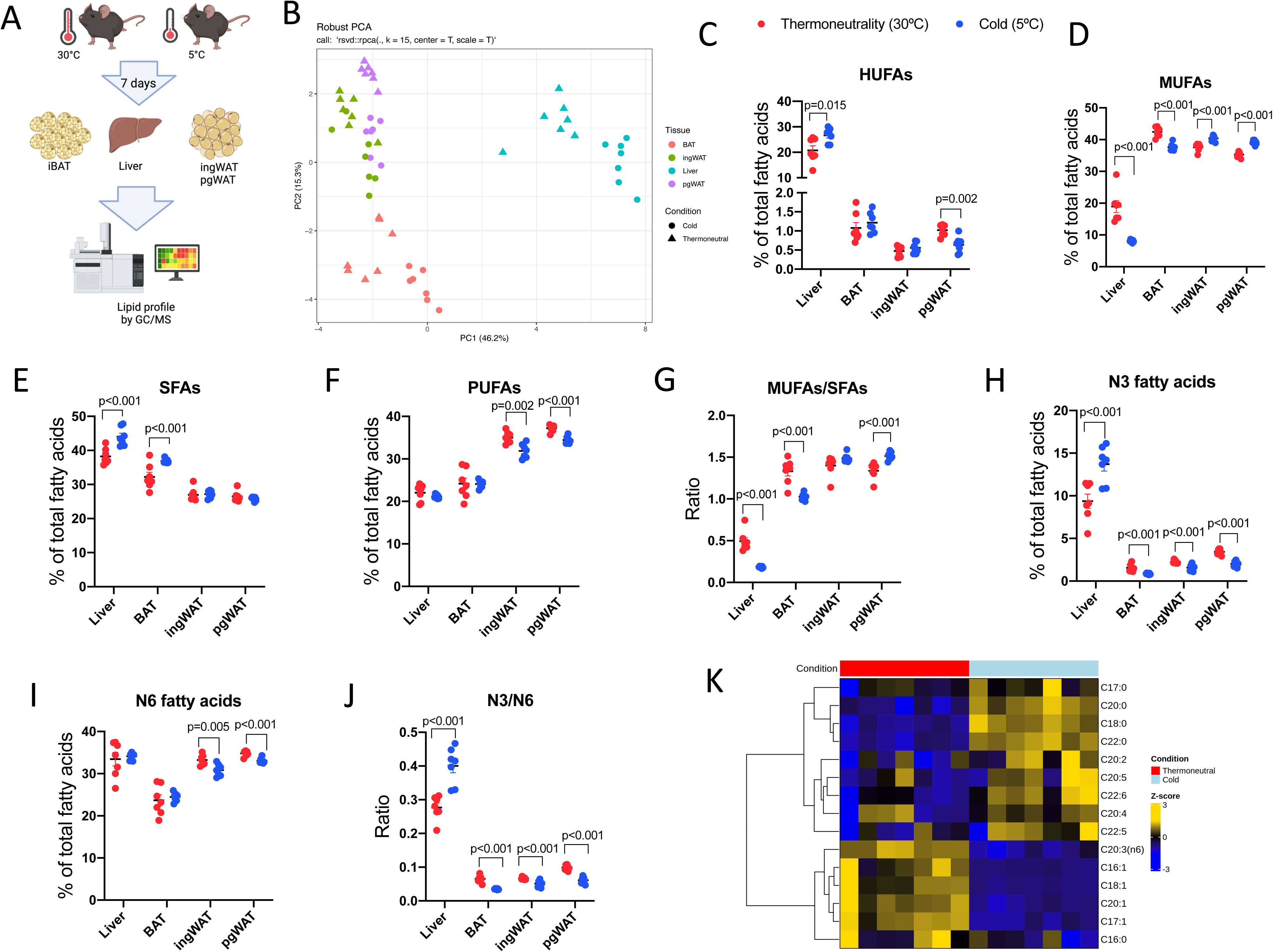
Tissue-specific lipid remodeling across adipose depots and liver upon cold exposure. **A** Schematic representation of the experimental design. Mice were exposed to cold (5 °C) or thermoneutrality (30 °C) for 7 days. Brown adipose tissue (BAT), inguinal white adipose tissue (ingWAT), perigonadal white adipose tissue (pgWAT), and liver were collected for fatty acid profiling by gas chromatography to assess depot-specific lipid remodeling. **B** Principal component analysis (PCA) of fatty acid composition across BAT, ingWAT, pgWAT and liver after 7 days of cold exposure. Each dot represents one biological replicate. Tissues segregate primarily by depot identity, indicating distinct lipid signatures, with cold-induced remodeling contributing to inter-tissue variance along the principal components. **C-F** Relative abundance of major fatty acid classes across tissues following cold exposure, highlighting differential enrichment of highly unsaturated (HUFAs), monounsaturated (MUFA) and saturated (SFA) and polyunsaturated fatty acids (PUFAs) between thermogenic BAT, subcutaneous ingWAT, visceral pgWAT, and liver. **G**, Index of SCD1-dependent desaturation (MUFAs/SFAs), indicating stearoyl-CoA desaturase activity across tissues in 5 °C and 30 °C. **H-I**, Quantification of total n-3 and n-6 fatty acids across the tissues in 5 °C and 30 °C. **J**, Scatter plot with the quantification of n-3/n-6 ratio in 5 °C and 30 °C. Data are expressed as mean ± S.E.M. n= 7 mice per group. Two-tailed unpaired Student’s t-test. **K**, Integrated heat map summarizing fatty acid abundance changes in the liver following cold exposure. *P*-values obtained by moderate t-test (limma). Only lipids with *p*≤0.05 are shown.

A more detailed analysis of cold-responsive lipid species in the liver revealed that, among the enriched HUFAs, three belonged to the n-3 class, namely docosahexaenoic acid (C22:6, DHA), eicosapentaenoic acid (C20:5, EPA), and docosapentaenoic acid (C22:5) (Fig. 1k). In contrast, arachidonic acid (C20:4, ARA) was the only upregulated n-6 lipid species in the liver (Fig. 1k). Interestingly, ingWAT and pgWAT displayed similar lipid changes, such as the increase in long chain MUFAs (C18:1, C20:1), and SFAs (C20:0) (Supplementary Fig. 1a-b). The modified lipid species in BAT also reflect a lower desaturation index in this tissue, especially demonstrated by the accumulation of stearic acid and reduction in oleic acid (Supplementary Fig. 1c). Collectively, these findings reveal that cold exposure drives a tissue-divergent lipid remodeling, thereby selectively reprogramming hepatic fatty acid composition, which is characterized by higher n-3 HUFAs content in contrast with a reduction in MUFAs levels.

### Chronic cold exposure selectively enriches hepatic n-3 HUFAs and suppresses fatty acid desaturation

The pronounced and distinct lipid remodeling observed in the liver in response to cold exposure prompted us to examine the temporal dynamics of hepatic lipid composition, enabling discrimination between acute stress-related changes and cold-acclimation–associated effects. To address this question, we designed a cold-exposure time-course experiment, in which mice were acclimated in a cold chamber and euthanized at multiple time points ranging from acute to chronic exposure (Fig. 2a). This time-course analysis revealed a transient accumulation of TGs within the first 24 hours of cold exposure, which resolved at later time points (from day 3 onward) (Fig. 2b). A similar temporal pattern was observed for the PUFA precursors alpha-linolenic acid (ALA, C18:3, n-3) and linoleic acid (LA, C18:2, n-6) (Fig. 2c–d). These findings suggest an initial hepatic burden of the most abundant PUFA precursors, likely resulting from the acute increase in adipose-to-liver lipolytic flux. However, with cold acclimation, these fatty acids are progressively metabolized. Interestingly, n-3 fatty acids increase after longer exposure (3 days), while n-6 content decreases (Fig. 2e). Consequently, the n-3/n-6 ratio increases upon chronic cold exposure (Fig. 2f) and coincides with the normalization of total TG content. An overall increase in hepatic HUFA content and HUFA/PUFA ratio also occurred after day 3 of cold exposure (Fig. 2g). While the precursor n-3 PUFA alpha-linolenic acid was transiently increased during the first days, DHA and EPA were found elevated at chronic time-points, consistent with the elevation of total n-3 HUFAs (Fig. 2h). Despite the acute fluctuations, the n-6 lipid species remained unchanged at the later time points of the time-course (Fig. 2h; Supplementary Fig. 2a–c). The exception was γ-linolenic acid (GLA, C18:3, n-6), which exhibited a consistent tendency to remain elevated during the subsequent stages of cold acclimation (Supplementary Fig. 2a). These findings suggest that HUFAs precursors are preferentially directed toward n-3 HUFA synthesis through elongation and desaturation reactions, rather than toward n-6–derived products in the liver. It is important to note that LA can also undergo alternative metabolic fates, including cytochrome P450 epoxygenase–mediated epoxidation followed by soluble epoxide hydrolase–dependent conversion to dihydroxy metabolites^12^. However, these pathways were not addressed in the present study.

**Figure 2.**
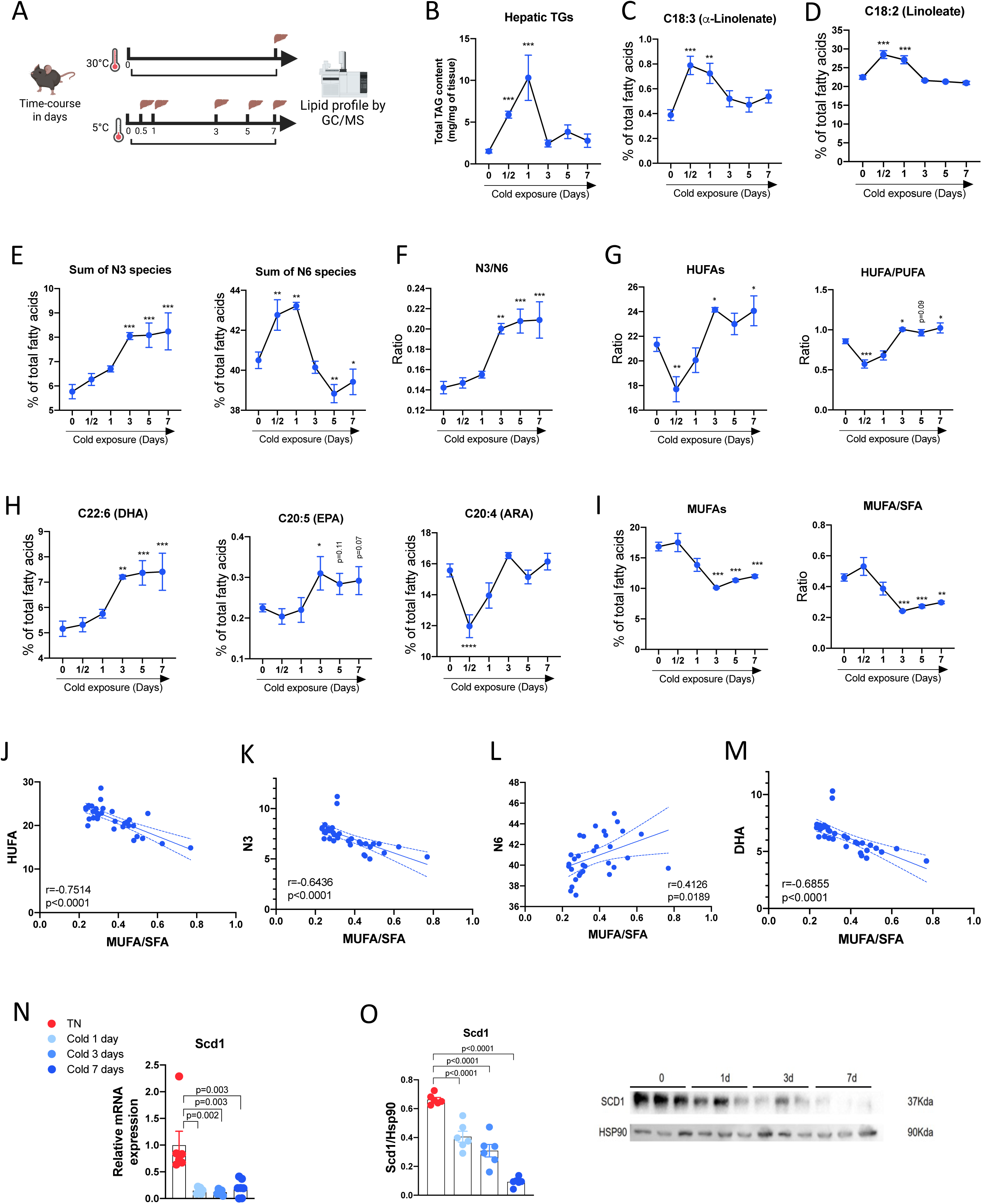
Time-course analysis of hepatic lipid remodeling during cold exposure. **A**, Schematic representation of the experimental design. Mice were exposed to cold for the indicated time points, and livers were collected along a defined time-course to assess dynamic metabolic adaptations induced by cold exposure. Gas chromatography coupled to mass spectrometry was used for profiling hepatic lipid remodeling over time. **B,** Hepatic triglyceride quantification during the cold exposure time-course. **C-D,** Quantification of α-linolenate (C18:3, n-3) and linoleate (C18:2, n-6) in the livers collected from mice during the cold exposure time-course. **E**, Fatty acid profiling of the sum of n-3 (B) and n-6 (C) over the time course in the liver. **F**, Quantification of n-3/n-6 ratio over the time-course. **G**, Quantification of total HUFAs and HUFAs/PUFAs ratio across the different time-points of cold time-course. **H**, Quantification of relative abundance of individual long-chain n-3 HUFAs (DHA and EPA) and the n-6 HUFA ARA, during cold time-course. **I**, Quantification of total MUFAs and MUFAs/SFAs ratio across the different time-points of cold time-course. Quantification of relative abundance of individual long-chain MUFAs and the Scd1-dependent desaturation index (MUFA/SFA) during cold time-course. Data are expressed as mean ± S.E.M. n= 4-6 mice per group. Differences were assessed using one-way ANOVA with false discovery rate control using the two-linear stet-up procedure of Benjamini, Krieger and Yekutieli. **J-M**, Pearson correlation between the SCD1-dependent desaturation index (MUFA/SFA) and the levels of total HUFAs (I), n-3 HUFAs (J), n-6 HUFAs (K) and DHA (L). The correlation coefficient (*r)* and corresponding two-tailed *P* value were calculated using Pearson’s method. **N**, mRNA expression of *Scd1* across time-points of 0 (30°C), 1, 3 and 7 days under 5°C. n=5-7 per group. **O**, Quantification and representative image of protein expression of SCD1 across time-points of 0 (30°C), 1, 3 and 7 days under 5°C. Data are expressed as mean ± S.E.M. n=6 mice per group. One-way ANOVA with false discovery rate control using the two-linear stet-up procedure of Benjamini, Krieger and Yekutieli.

Interestingly, the reduction in MUFA content and the associated decrease in the SCD1-dependent desaturation index were also exclusively observed upon cold acclimation (Fig. 2i; Supplementary Fig. 2d–f). SCD1-dependent desaturation index displayed inverse correlations with both total HUFAs and n-3 fatty acid levels, suggesting that these lipid species may directly modulate hepatic SCD1 activity during cold exposure (Fig. 2j–k). In contrast, n-6 fatty acid levels were positively associated with the desaturation index (Fig. 2l). Among the n-3 upregulated lipids, DHA exhibited the strongest inverse correlation with the desaturation index (Fig. 2m), although EPA also showed similar trends (Supplementary Fig. 2g). Consistent with these lipid profile changes, both mRNA and protein expression of SCD1, the enzyme responsible for desaturation of SFAs into MUFAs, were reduced in the liver upon cold acclimation (Fig. 2n–o). Other *de novo* lipogenesis (DNL) pathway markers were also reduced by chronic cold exposure at both gene and protein levels upon cold acclimation (Supplementary Fig. 3a–b). Livers from cold exposed mice displayed preserved mitochondrial oxidative phosphorylation (OXPHOS), despite the reduced β-oxidation capacity in response to palmitoyl-carnitine (Supplementary Fig. 3c-d), along with accumulation of long-chain acylcarnitines, consistent with limited entry of long-chain fatty acids into oxidative pathways (Supplementary Fig. 3e). Cold also enhanced the coenzyme-Q levels (Supplementary Fig. 3f), which may reflect a compensatory adaptation to sustain electron transport chain efficiency despite reduced fatty acid-derived substrate supply. Importantly, total cardiolipin content and citrate synthase activity remained unchanged, indicating preserved mitochondrial content (Supplementary Fig. 3g–h). Together, these observations indicate that hepatic lipid remodeling during cold adaptation occurs in a metabolic context where fatty-acid oxidation is relatively limited. Accordingly, the changes in lipid composition are unlikely to reflect increased oxidative metabolism but instead suggest altered desaturation dynamics and coordinated suppression of de novo lipogenesis (DNL). Consistent with this, previous work shows that acute cold exposure reduces fatty-acid contribution to the TCA cycle while increasing reliance on glucose and branched-chain amino acids^14^.

### Increase in n-3 HUFAs suppresses SCD1 activity in hepatocytes

The inverse correlation between n-3 HUFAs and the MUFA/SFA ratio observed in the liver from cold-exposed mice prompted us to examine a possible causality between the increase of n-3 HUFAs and the reduction in SCD1-dependent desaturation index. As a first strategy, we used *fat-1* transgenic mice, which express the *C. elegans* desaturase *fat1*, enabling endogenous conversion of n-6 into n-3 fatty acids, thereby increasing n-3/n-6 ratio regardless of the thermal stress^22^ (Fig. 3a). As expected, we found higher levels of n-3 fatty acids, while total n-6 fatty acid content remained unchanged (Fig. 3b). In line with this, n-3/n-6 ratio was found higher in the liver of *fat-1* mice, which confirmed the *fat1* gene activity in these transgenic animals (Fig. 3c). DHA, EPA, and DPA levels were elevated in *fat-*1 mice (Fig. 3d). Notably, *fat-1* mice exhibited a reduced SCD1-dependent desaturation index (Fig. 3e), which negatively correlated with n-3 but not with n-6 fatty acid levels (Fig. 3f–g). This inverse association was similarly observed across individual n-3 species, including DHA, EPA, DPA, and ALA (Fig. 3h–k). Together, these data suggest that the increase in n-3 HUFAs leads to the reduction in hepatic desaturation index, as selective enrichment of n-3 species alone in the *fat-1* mouse model was sufficient to lower SCD1-dependent desaturation activity. To determine whether n-3 HUFAs directly regulate *Scd1* expression or whether the reduction observed *in vivo* results from increased sympathetic activity during cold exposure, AML12 hepatocytes were treated with DHA or norepinephrine for 48 h. DHA drastically reduced *Scd1* mRNA expression, whereas norepinephrine had no effect (Fig. 3l–m), indicating that the decrease in hepatic *Scd1* observed *in vivo* is driven by the action of n-3 fatty acids rather than by cold-induced sympathetic activation.

**Figure 3.**
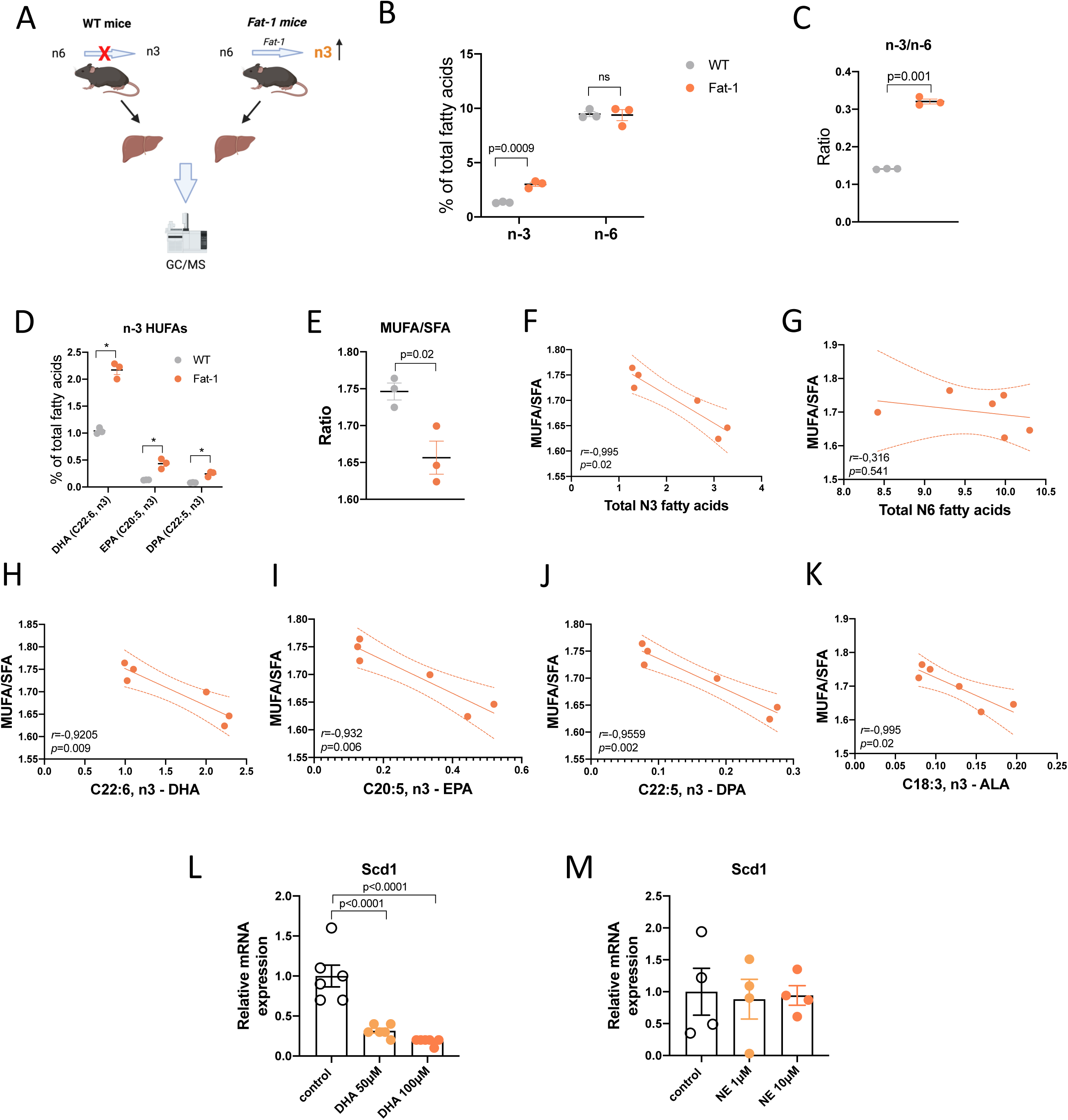
Livers from *Fat-1* mice recapitulate lipid profile of cold acclimated mice. **A**, Schematic representation of the experimental design. Livers from wild-type and *Fat-1* mice kept under room temperature were harvested and utilized for lipid profiling through gas chromatography coupled to mass spectrometry. *Fat-*1mice express a desaturase that converts n-6 into n-3 fatty acids, thus artificially increasing n-3/n-6 ratio. **B**, Fatty acid profiling of the sum of n-3 and n-6 fatty acids in the liver of both WT and *Fat-*1mice. **C**, Quantification of n-3/n-6 ratio in the liver of both WT and *Fat-*1mice. **D**, Quantification of n-3 HUFAs species in the liver of both WT and *Fat-*1mice. **E**, Quantification of MUFAs/SFAs ratio in the liver of both WT and *Fat-*1mice. Data are expressed as mean ± S.E.M. n=3 mice per group. Two-tailed unpaired Student’s *t*-test. **F-K**, Pearson correlation between the SCD1-dependent desaturation index (MUFA/SFA) and the levels of total n-3 FAs (F), n-6 FAs (G), DHA (H), EPA (I), DPA (J) and ALA (K). The correlation coefficient (*r)* and corresponding two-tailed *P* value were calculated using Pearson’s method. N=6 mice. **L**-**M**, mRNA expression of Scd1 in AML12 hepatocytes following treatment with docosahexaenoic acid (DHA; 50 or 100 μM) for 48 h (L) or norepinephrine (NE; 1 or 10 μM) for 48 h (M). Data are expressed as mean ± S.E.M. n= 4-6 biological replicates. One-way ANOVA with false discovery rate control using the two-linear stet-up procedure of Benjamini, Krieger and Yekutieli.

We also tested whether this negative association between n-3 HUFAs and the desaturation index is maintained in a steatosis condition, in which a higher desaturation index is expected to happen. To this end, we quantified hepatic lipid species in mice fed a high-fat diet (HFD) or HFD supplemented with oral fructose (Supplementary Fig. 4a), which represent models of diet-induced obesity associated with hepatic steatosis and metabolic steatohepatitis (MASH), respectively. Both HFD and HFD + fructose mice exhibited reduced hepatic n-3 fatty acid content and a lower n-3/n-6 ratio (Supplementary Fig. 4b–c). All major n-3 HUFAs were decreased in these models, with DHA — the most abundant n-3 HUFA —showing the most pronounced reduction (Supplementary Fig. 4d). Consistent with these changes, the SCD1-dependent desaturation index was elevated in both HFD and HFD + fructose mice, whereas increased *Scd1* mRNA expression was observed only in the MASH model (Supplementary Fig. 4e–f). In agreement with findings from cold-exposed mice, both total n-3 and n-6 levels exhibited inverse associations with the desaturation index (Supplementary Fig. 4g). At the individual lipid level, either the n-3 or the n-6 HUFAs negatively associated with desaturation index (Supplementary Fig. 4h).

### Cold adaptation engages HUFA biosynthesis machinery in both adipose tissue and liver

Having established that the higher hepatic n-3 HUFAs content is strongly associated with reduced SCD1-dependent desaturation in the liver of cold acclimated mice, we next sought to investigate the potential origin of these fatty acids. Specifically, we asked whether the observed changes in HUFA abundance in the liver reflect alterations in the expression of *Fads1* and 2 in liver or adipose depots, key enzymes that encode the expression of Δ5- and Δ6-desaturases, respectively, which mediate the conversion of essential fatty acids into long-chain HUFAs. The biosynthetic pathways for n-3 and n-6 HUFAs, operating in parallel and sharing the same desaturation and elongation machinery, are schematically depicted in Fig. 4a. We sought to assess *Fads1* and *Fads2* expression to gain insight into whether hepatic HUFA availability is primarily driven by local biosynthesis or influenced by lipid flux from adipose tissue.

**Figure 4.**
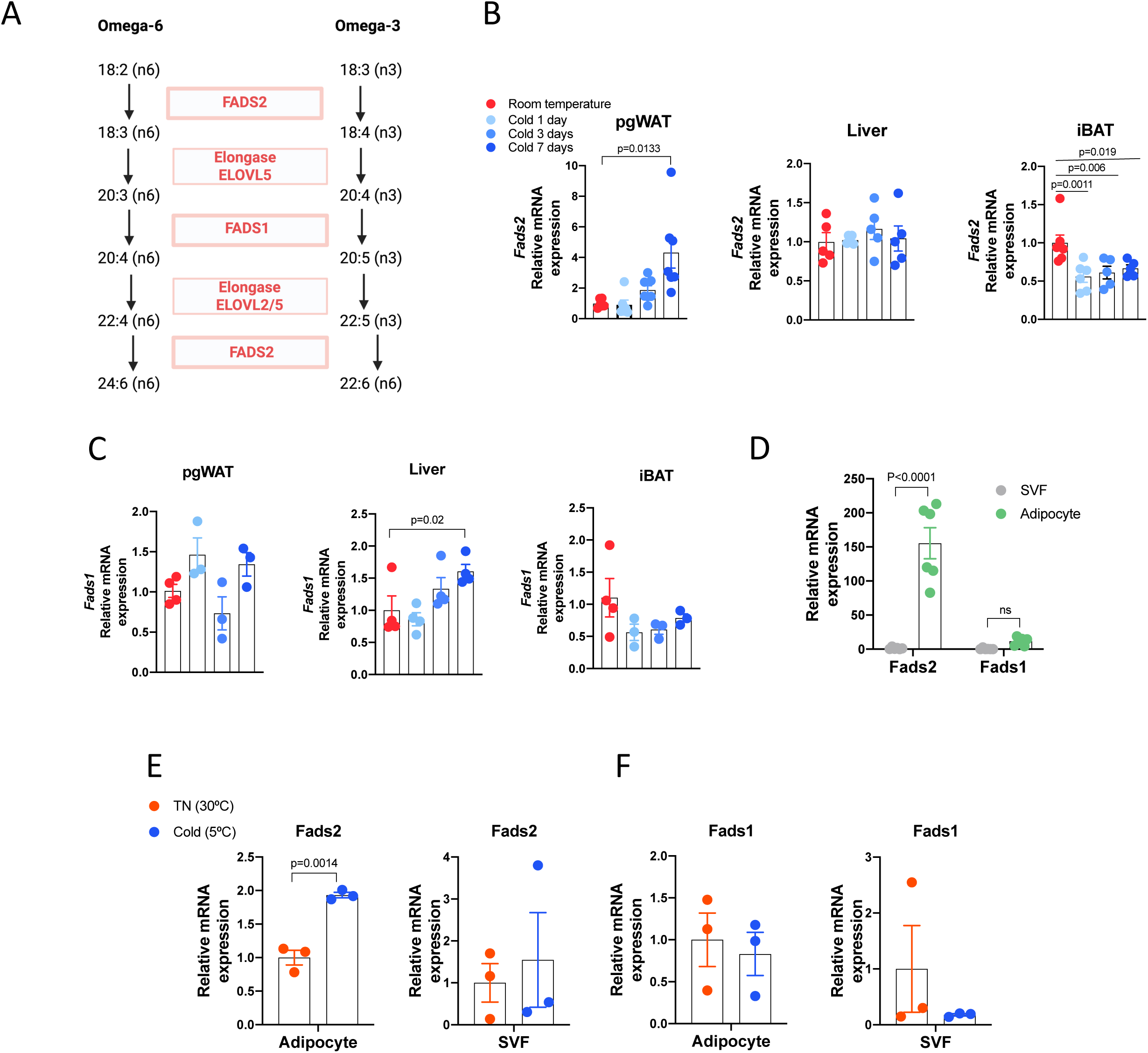
Involvement of fatty acid desaturases 1 and 2 in the cold-induced increase of hepatic HUFAs. **A**, PUFAs’elongation and desaturation cascade for the biosynthesis of HUFAs. N-3 and n-6 PUFAs are converted through successive and coordinated elongation and desaturation steps into HUFAs. Elovl 2 and 5 are the elongases, while Fads1 and 2 are the desaturases acting in this cascade. **B-C**, mRNA expression of *Fads2*(B) *and Fads1*(C) in BAT, pgWAT and liver across time-points of 0 (30 °C), 1, 3 and 7 days under 5 °C. n=3-7 mice per group. One-way ANOVA with false discovery rate control using the two-linear stet-up procedure of Benjamini, Krieger and Yekutieli. **D**, mRNA expression of *Fads2 and Fads1* in stromal vascular fraction (SVF) and adipocyte fraction of pgWAT from C57BL6/J mice. n=6 mice per group. **E-F**, mRNA expression of *Fads2* (E) *and Fads1* (F) in SVF and adipocyte fraction of pgWAT from C57BL6/J mice exposed to 5 °C or 30 °C for 7 days. n=3 mice per group. Data are expressed as mean ± S.E.M. Two-tailed unpaired Student’s *t*-test.

We observed a significant induction of Fads2 expression at both the mRNA and protein levels across WAT depots, whereas BAT and liver did not exhibit a comparable increase (Fig. 4b, Supplementary Fig. 5a–c). In contrast, *Fads1* mRNA expression was selectively upregulated in the liver in response to cold exposure, without changes in BAT or pgWAT (Fig. 4c). Consistent with enhanced hepatic desaturation capacity, cold exposure increased the ratios of HUFAs to PUFAs, as well as product-to-precursor indices including DHA/ALA (C18:3, n3), EPA/ALA (C18:3, n3), and ARA/LA (C18:2, n6) (Supplementary Fig. 5d–g), supporting increased flux through the HUFA biosynthetic pathway in the liver. To determine whether *Fads2* expression within white adipose tissue originates predominantly from adipocytes or other cellular components of the adipose niche, we isolated mature adipocytes and the stromal vascular fraction (SVF) from pgWAT. *Fads2* mRNA levels were markedly higher in adipocytes compared to the SVF, whereas *Fads1* expression was comparable between these fractions (Fig. 4d). Notably, cold exposure selectively increased *Fads2* expression in the adipocyte fraction, while *Fads1* remained unchanged (Fig. 4e–f). To determine the cellular compartment in which fatty acid desaturases are expressed in the liver, we analyzed single-cell RNA sequencing data from the publicly available Liver Cell Atlas (https://www.livercellatlas.org). This analysis revealed that both *Fads1* and *Fads*2 are predominantly expressed in hepatocytes (Supplementary Fig. 5g). This suggests that the increased hepatic HUFA levels observed upon cold exposure may reflect contributions from both adipose-derived fatty acids and liver-intrinsic biosynthesis.

### Cold exposure enriches hepatic triglycerides with n-3 HUFAs while depleting MUFA-containing species

Having established that n-3 HUFAs accumulate in the liver upon cold exposure, we next sought to determine the lipid species in which these fatty acids are incorporated. To this end, we performed an untargeted lipidomic analysis (LC–MS/MS) of livers from mice exposed to thermoneutrality or cold for 7 days (Fig. 5a). Principal component analysis revealed a clear segregation between groups, indicating extensive remodeling of the hepatic lipidome (Fig. 5b). To identify the lipid classes contributing to this separation, we next examined individual lipid species and lipid classes. This analysis revealed that TGs and FFA recapitulated the changes previously observed through GC/MS, displaying marked remodeling characterized by enrichment of HUFA-containing species alongside a reduction in MUFAs (Fig. 5c–g), resulting in a lower desaturation in both TG and FFA forms (Fig. 5f–g). As a result, we observed a negative correlation between n-3/n-6 HUFAs ratio and MUFA/SFA ratio (Fig. 5h). In this context, enrichment of n-3 HUFAs within the free pool further supports suppression of hepatic SCD1 activity, thereby stabilizing this lipid compositional state. The sum of either SFA or PUFAs was not modified by cold (Supplementary Fig. 6a–b). When narrowing down the analysis through the individual species, we found consistent upregulation of HUFAs and downregulation of MUFAs in both FFA and TG forms (Fig. 5i–j). In line with these findings, heatmap visualization showed that most of TG species upregulated by cold contained DHA within their acyl backbone (Fig. 5k). Moreover, some PUFAs in the form of FFAs were reduced in the liver of cold-exposed mice (Supplementary Fig. 6c). Consistently, the HUFA/PUFA ratio was elevated by cold, corroborating the data we observed through gas chromatography, once more highlighting the increased FADS-mediated desaturation process in the liver (Supplementary Fig. 6d–e).

**Figure 5.**
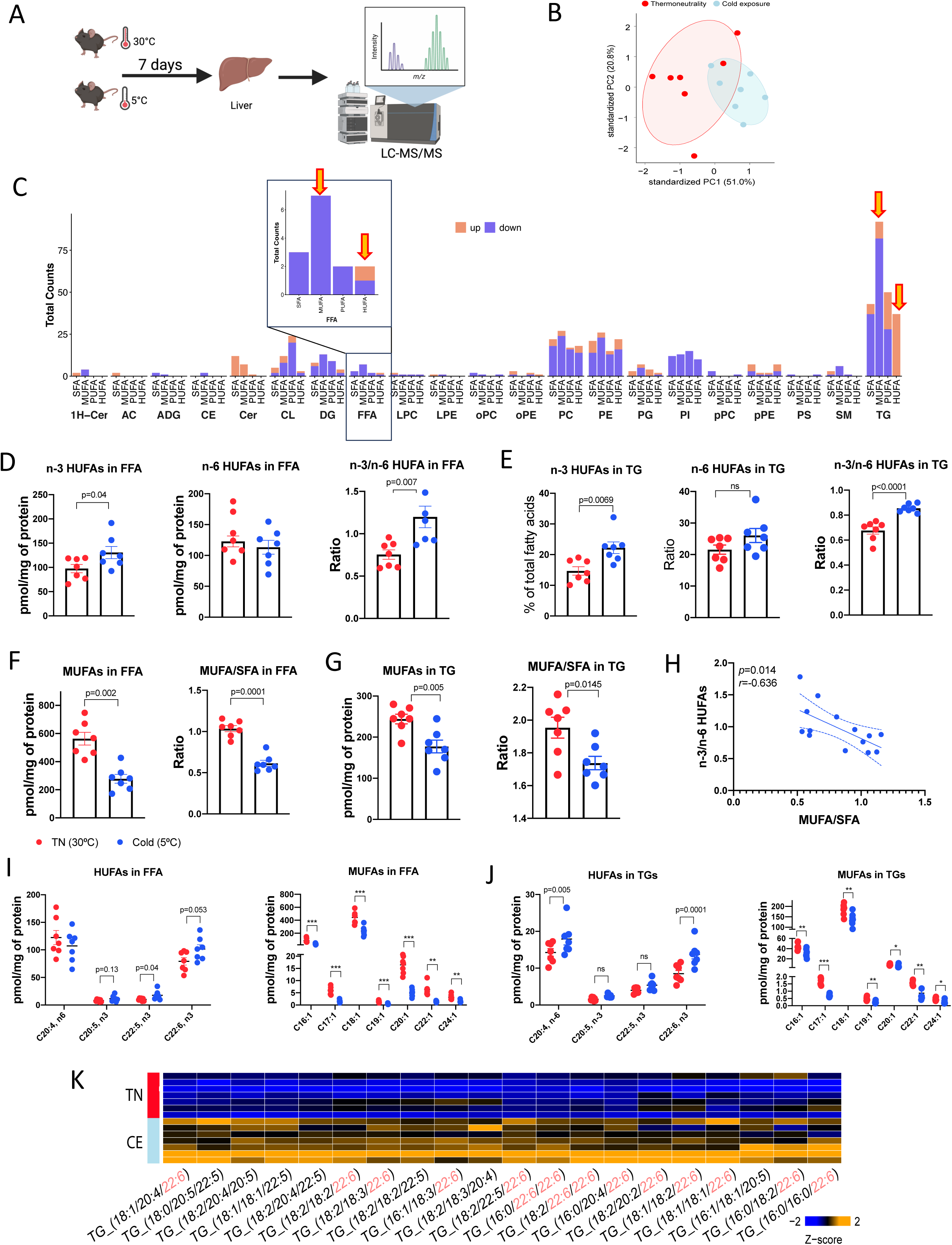
Incorporation of HUFAs into FFA and TGs in the liver of cold-exposed mice. **A**, Schematic representation of the experimental design. Mice were exposed to 5 °C or 30 °C for 7 days. Then, liver was collected for untargeted lipidomic analysis through liquid chromatography-tandem mass spectrometry (LC-MS/MS). Highlighted are the similar changes in HUFAs and MUFAs in the free fatty acid (FFA) and triglyceride (TG) forms. **B**, PCA of fatty acid composition in the liver from mice exposed to 5 °C or 30 °C for 7 days. Each dot represents one biological replicate. **C**, Quantification of up- and down-regulated lipid classes (MUFAs, PUFAs, SFAs and HUFAs) by cold, across all the detected lipid species in the liver. Highlighted are the similar changes in HUFAs and MUFAs in the free fatty acid (FFA) and triglyceride (TG) forms. **D**, Quantification of total n-3 and n-6 HUFAs as well as the n-3/n-6 ratio in the FFA form in the liver of mice exposed to 5 °C or 30 °C. **E**, Quantification of total n-3 and n-6 HUFAs as well as the n-3/n-6 ratio in the TG form in the liver of mice exposed to 5 °C or 30 °C. **F**, Quantification of total MUFAs as well as the MUFA/SFA ratio in the FFA form in the liver of mice exposed to 5 °C or 30 °C. **G**, Quantification of total MUFAs as well as the MUFA/SFA ratio in the TG form in the liver of mice exposed to 5 °C or 30 °C. **H**, Correlation between the SCD1-dependent desaturation index (MUFA/SFA) and the n-3/n-6 HUFAs ratio. The correlation coefficient (*r)* and corresponding two-tailed *P* value were calculated using Pearson’s method. n=12 mice per group. **I**, Quantification of individual species of HUFAs and MUFAs as FFA in the liver of mice exposed to 5 °C or 30 °C. **J,** Quantification of individual species of HUFAs and MUFAs as TGs in the liver of mice exposed to 5 °C or 30 °C. **K**, Integrated heat map summarizing the significantly cold-induced TGs in the liver following cold exposure. Data are expressed as mean ± S.E.M. n= 7 mice. Two-tailed unpaired Student’s *t*-test.

These results indicate that cold acclimation reprograms lipid composition upstream of TG synthesis and assembly in hepatocytes. The presence of this compositional signature in the FFA pool suggests that altered fatty acid availability and desaturation, rather than selective esterification, contribute to the enrichment of HUFAs within hepatic TGs. Consistent with a highly dynamic hepatic lipid state that permits TG remodeling, cold exposure also increased the protein abundance of key enzymes involved in both TG lipolysis and re-esterification, including adipocyte triglyceride lipase (ATGL), hormone-sensitive lipase (HSL), and cytosolic phosphoenolpyruvate carboxykinase (PEPCK-C), while we found no changes in glycerol kinase (GK) expression (Supplementary Fig. 6f).

### Cold-induced hepatic lipid remodeling is propagated to circulating VLDL particles

Given that hepatic TGs constitute the primary lipid pool exported from the liver in the form of VLDL, we next asked whether the TG remodeling observed upon cold exposure extends to circulating lipoproteins. To address this, we performed an untargeted LC–MS/MS lipidomic analysis of the VLDL/IDL fraction isolated from plasma (Fig. 6a). This approach allowed us to assess whether the cold-induced adipose–liver lipid axis influences hepatic lipid export and, consequently, the distribution of lipid species to peripheral tissues via lipoprotein-mediated transport. TGs represented the most diverse lipid class, consistent with the structural heterogeneity of neutral lipid species within the lipoprotein core (Supplementary Fig. 7a). In terms of abundance, phosphatidylcholines and cholesteryl esters were the predominant lipid classes detected, followed by TGs, phosphatidylethanolamines, free cholesterol and others (Supplementary Fig. 7b).

**Figure 6.**
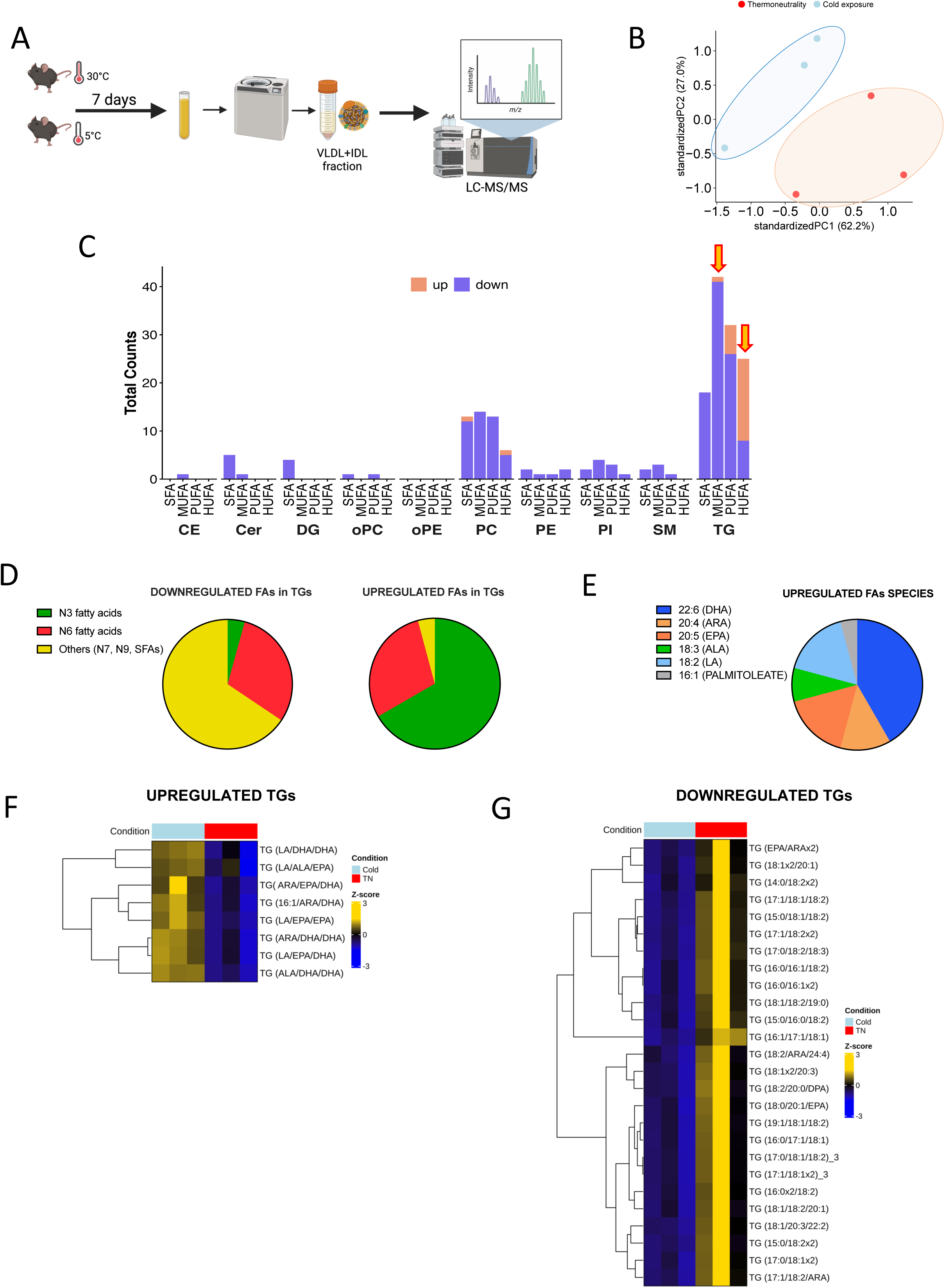
Cold-induced incorporation of HUFAs into TGs in circulating VLDL/IDL fractions. **A**, Schematic representation of the experimental design. Mice were exposed to 5 °C or 30 °C for 7 days. Then, blood was collected, and plasma fraction used for obtaining the VLDL/IDL fractions for untargeted lipidomic analysis through LC-MS/MS. **B**, PCA of of the VLDL/IDL lipidome from mice exposed to 5 °C or 30 °C for 7 days. Each dot represents one biological replicate. **C**, Quantification of up- and down-regulated lipid classes (MUFAs, PUFAs, SFAs and HUFAs) by cold, across all the detected lipid species in VLDL/IDL fraction. Highlighted are the similar changes in HUFAs and MUFAs in the FFA and TG forms. **D**, Pizza graphs illustrating the proportion of significantly down- and up-regulated n-3 and n-6 fatty acids incorporated into TGs in VLDL/IDL fraction. **E,** Pizza graphs illustrating the proportion of significantly up-regulated individual n-3 and n-6 lipid species into TGs in VLDL/IDL fraction. **F**, Integrated heat map summarizing the significantly cold-induced TGs in the VLDL/IDL fraction following cold exposure. **G**, Integrated heat map summarizing the significantly cold-suppressed TGs in the VLDL/IDL fraction following cold exposure. n= 3 mice. N=7 mice per group. Lipids shown have a nominal *p* value ≤0.05, determined by moderate t-statistics.

Principal component analysis of the VLDL/IDL lipidome revealed a clear separation between thermoneutral and cold-exposed groups, indicating significant remodeling of circulating lipoproteins in response to cold (Fig. 6b). Likewise observed in the liver, TGs within the VLDL/IDL fraction exhibited marked remodeling, characterized by enrichment of HUFA-containing TG species accompanied by a reduction in MUFA-containing TGs (Fig. 6c). Importantly, most of the increased HUFA-containing TG species corresponded to n-3 HUFAs, with DHA representing the most abundant lipid specie (Fig. 6d–e). The heat map in Fig. 6f indicates that the majority of upregulated TG species in the VLDL/IDL fraction were enriched in DHA-containing species. In contrast, the heat map in Fig. 6g illustrated that most of the reduced TG species were predominantly composed of MUFAs. Together, these findings indicate that cold-induced hepatic TG remodeling is propagated to circulating lipoproteins, resulting in preferential export of n-3 HUFA–enriched TG species and reshaping systemic lipid distribution.

## Discussion

The present study was motivated by a central metabolic paradox regarding why increased adipose-to-liver lipid flux leads to sustained steatosis in obesity, whereas during cold exposure hepatic lipid accumulation is only transient and resolves with acclimation despite persistently elevated flux. Our findings support the model that the outcome of lipid influx is not dictated by flux magnitude alone, but by the qualitative nature of the lipid species being delivered, processed, and distributed systemically. By combining lipid profiling through GC/MS, temporal analyses, and untargeted lipidomics of hepatic and circulating lipoproteins, we demonstrate that cold acclimation induces a coordinated remodeling of hepatic lipid composition characterized by enrichment of n-3 HUFAs, which drive the suppression of DNL and concomitant depletion of MUFAs through the suppression of SCD1-dependent desaturation. The MUFA-to-HUFA lipid profile shift occurs across both FFA and TG pools in the liver, indicating that compositional rewiring precedes and shapes lipid storage and export. Importantly, this remodeled hepatic lipid profile is propagated to the circulation via VLDL and IDL particles, in which n-3 HUFA–containing TGs replace MUFA-enriched species (Fig. 7). Overall, by increasing endogenous n-3-HUFAs availability, the liver transcriptionally and functionally suppresses DNL and SCD1-dependent desaturation, thereby limiting MUFA-driven TG assembly and buffering hepatic lipid accumulation despite elevated fatty acid influx resulting from adipose tissue lipolysis.

**Figure 7.**
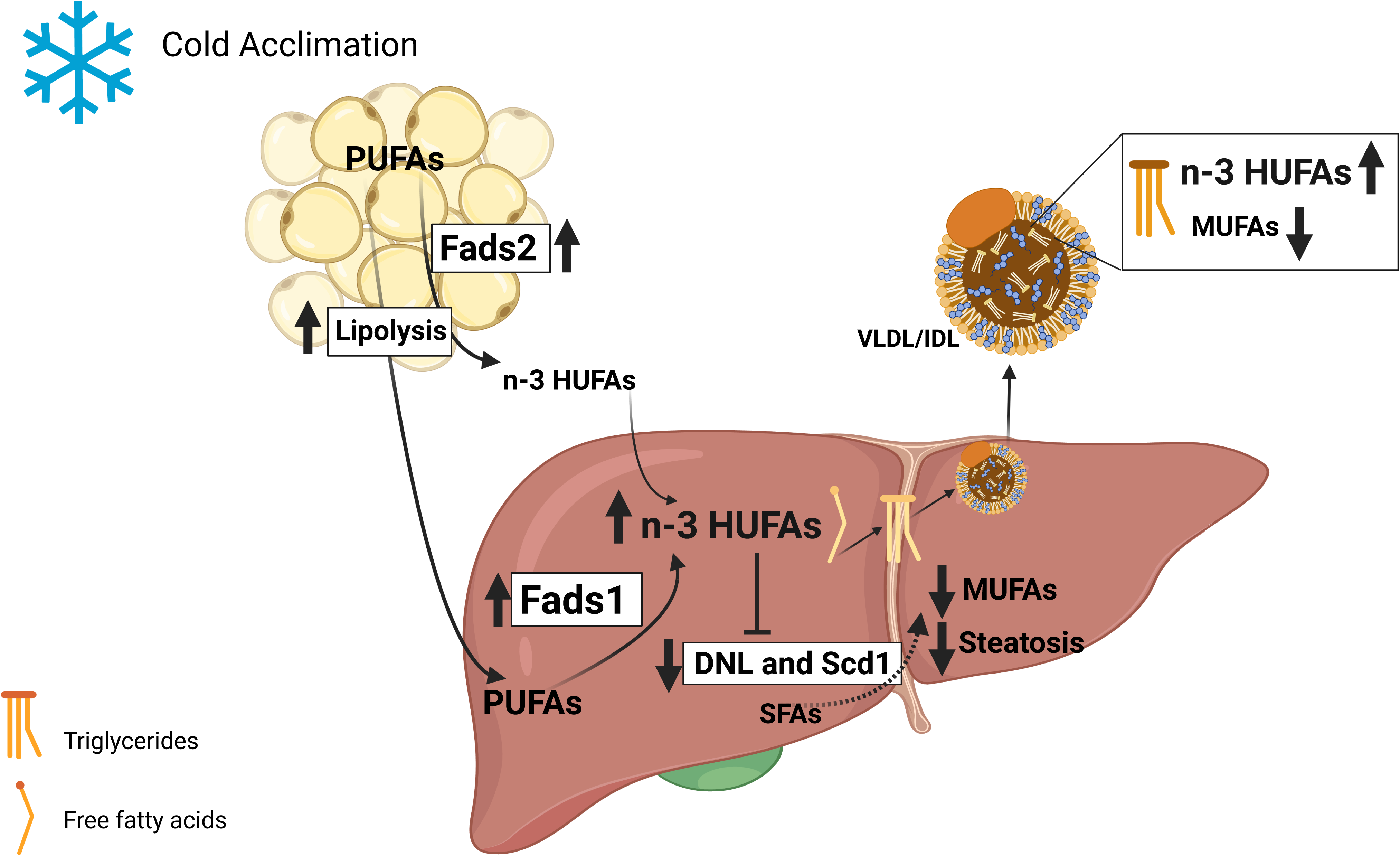
Graphical abstract summarizing the main findings of the study. Cold acclimation promotes the mobilization of PUFAs from adipose tissue (AT), which are released directly or converted locally into n-3 HUFAs via FADS1/2-mediated desaturation. These fatty acids are delivered to the liver, where additional conversion of PUFAs to n-3 HUFAs further enhances hepatic n-3 HUFA levels. Elevated hepatic n-3 HUFAs suppress *de novo* lipogenesis (DNL) and SCD1-dependent desaturation, reducing MUFA availability for triglyceride (TG) synthesis and limiting steatosis. Concurrently, n-3 HUFA–enriched TGs are assembled and exported as VLDL particles, resulting in a remodeled circulating lipoprotein profile characterized by high n-3 HUFA and low MUFA content.

An important insight from our study is that the coordinated regulation of FADS1 and 2 activity/expression during cold adaptation is associated with the suppression of *de novo* lipogenesis in the liver, especially SCD1-dependent desaturation, which in turn is required for hepatic TG synthesis^23–25^. We observed upregulation of *Fads2* predominantly in white adipose tissue and induction of *Fads1* in the liver. Desaturase activity is regulated not only at the transcriptional level but also through substrate availability, post-transcriptional mechanisms, and metabolic context. Supporting this notion, we detected increased HUFA/PUFA ratios and elevated product-to-precursor ratios (e.g., DHA/ALA), consistent with enhanced FADS activity in the liver. The functional relevance of *Fads1 and 2* activities for the suppression of hepatic DNL is supported by prior studies showing that global loss of either *Fads1* or *Fads2* leads to hepatic steatosis and dyslipidemia^26,27^. MacLeod *et al.*^26^ demonstrated that ALA needs to be converted into EPA and DHA by *Fads* in order to suppress hepatic lipogenesis. Indeed, the capacity of long-chain HUFAs to suppress DNL and *Scd1* is well documented through both *in vivo* and *in vitro* studies^19,26,28^.

It is noteworthy that similar lipid remodeling has been observed at the TG level under fasting^29^, a physiological condition that likewise increases adipose tissue lipolysis and hepatic fatty acid influx^30^. Sixteen hours of fasting has been reported to enrich hepatic TGs in n-3 HUFAs while reducing MUFA content in mice^29^, closely resembling the compositional shift described here. Rather than representing a cold-exclusive mechanism, enhanced HUFA biosynthesis may constitute a generalized metabolic safeguard engaged whenever hepatic lipid influx exceeds oxidative capacity. Importantly, this stands in sharp contrast to obesity, where chronic adipose tissue lipolysis likewise drives hepatic fatty acid influx but leads to elevated MUFAs synthesis, increased SCD1 activity, and progressive TG accumulation^31,32^. Thus, it is not lipid flux per se that dictates hepatic outcome, but the type of fatty acids and their enzymatic handling in the liver or adipose tissue. In the setting of cold adaptation, we showed that enhanced conversion of PUFAs into long-chain n-3-HUFAs by FADS is a key step for the suppression of DNL and SCD1-driven TG assembly, thereby buffering hepatic lipid overload despite persistent adipose-derived fatty acid flux.

Despite the fact that both cold exposure and obesity enhance adipose-to-liver lipid flux, a key distinguishing feature between these conditions is the hepatic n-3/n-6 PUFA ratio. Previous studies have reported that MASLD/MASH are characterized by a reduced n-3/n-6 ratio accompanied by increased SCD1 activity^31,33,34^. In contrast, our findings demonstrate that cold acclimation induces the opposite phenotype, which contributes to protection against hepatic steatosis. Δ5- and Δ6-desaturases, encoded by *Fads1* and *Fads2*, respectively, catalyze key desaturation steps in both the n-3 and n-6 PUFA pathways. However, it remains unclear whether, and by which mechanisms, these enzymes preferentially channel substrates toward the n-3 rather than the n-6 branch. Our data raise the possibility that such selective partitioning may occur during cold exposure compared with thermoneutral conditions. In obesity, the shift toward the n-6 cascade may be more readily explained by reduced dietary availability of n-3 species, resulting in greater relative abundance of n-6 substrates for enzymatic conversion. By contrast, the mechanisms governing Δ5- and Δ6-desaturase activity toward either PUFA cascade under distinct physiological stimuli, such as cold exposure, remain poorly understood. Elucidating how these enzymes are differentially regulated across metabolic contexts represents an important frontier for future investigation.

Using *fat-1* transgenic mice, which endogenously convert n-6 to n-3 PUFAs, we increased the n-3/n-6 ratio independently of cold exposure. Remarkably, *fat-1* mice phenocopied cold-exposed mice especially regarding the inverse correlation between n-3/n-6 ratio and SCD1 activity. Importantly, these mice are known to be protected against HFD-induced hepatic steatosis^32^. These findings suggest that modulation of the n-3/n-6 ratio is not merely associative but is mechanistically sufficient to drive the protective hepatic phenotype induced by cold.

Mechanistically, the inverse relationship between n-3 HUFAs and SCD1 activity observed in the liver of cold-acclimated mice is supported by foundational work identifying a PUFA-responsive element within the *Scd1* promoter, which mediates direct transcriptional repression by long-chain HUFAs, including ARA and EPA^35^. Beyond promoter-level regulation, n-3 HUFAs such as DHA suppress hepatic lipogenic programs through membrane receptor-dependent mechanisms. In primary hepatocytes, DHA reduced SREBP-1c abundance and downstream lipogenic genes, including *Fasn* and *Scd1*, in a GPR40 (FFAR1)-dependent manner, an effect reversed by pharmacological antagonism of GPR40^36^. Conversely, dietary n-3 HUFA deficiency increases hepatic SCD1 expression and desaturation indices, including the MUFA/SFA ratio used here, in rodent models^37^. Collectively, these studies support our mechanistic model in which enhanced conversion of precursor PUFAs into long-chain HUFAs during cold adaptation serves not merely as a compositional shift but as an active regulatory program.

The role of SCD1 in hepatic lipid homeostasis is complex and highly context dependent. Genetic manipulation studies have demonstrated that altered SCD1 activity profoundly impacts hepatic lipid balance. Whole-body SCD1 deficiency protects against diet-induced obesity and hepatic triglyceride accumulation, largely by limiting MUFA synthesis and de novo lipogenesis^38,39^. In contrast, liver-specific disruption of SCD1 activity can lead to accumulation of saturated fatty acids, endoplasmic reticulum stress, inflammation, and, in some settings, exacerbation of liver injury despite reduced TG storage^40,41^. In this context, the concomitant enrichment of n-3 HUFAs during cold adaptation may mitigate the lipotoxic effects associated with SFA accumulation resulting from reduced MUFA synthesis.

We also show that cold exposure propagates hepatic lipid remodeling to the circulation by selectively enriching VLDL/IDL triglycerides with n-3 HUFAs at the expense of MUFA-containing species. It was recently demonstrated that HUFAs are preferentially taken up by hepatocytes in culture and further incorporated into TGs, when DNL is suppressed^42^. This goes in close agreement with our data, as we found that n3-HUFAs incorporation into TGs coincides with an overall decrease in gene and protein expression of DNL enzymes. Thus, it is likely that the increased availability of n-3-HUFAs as FFAs in hepatocytes under cold, causes the suppression of the DNL and then the incorporation of these lipids into TGs. Given that hepatic TGs are secreted as VLDL, changes in hepatic fatty acid composition are likely transmitted systemically through lipoprotein remodeling. Consistent with this model, previous study demonstrated that acute cold exposure increases circulating HUFA-containing TGs in both humans and mice, an effect attributed to their incorporation into VLDL particles^21^. This shift in lipoprotein lipid quality is likely to alter both lipoprotein–tissue interactions and downstream remodeling into LDL particles. Beyond serving as a delivery system for bioactive fatty acids to peripheral tissues, n-3-HUFA enriched VLDL particles may themselves represent a less atherogenic lipoprotein species, consistent with epidemiological and experimental evidence linking n-3 HUFA–rich lipid profiles to cardiovascular protection^43–45^. Moreover, it is known that fatty acids delivery by TG rich lipoproteins, like VLDL and IDL, into thermogenic fat is essential for cold adaptation^5,46^. In BAT, binding of VLDL to the VLDL receptor (VLDLR) promotes uptake of TG-derived fatty acids, providing the fuel necessary to maintain thermogenic function^46^. Accordingly, VLDLR-deficient mice exhibit impaired cold tolerance^46^. Thus, cold-induced reprogramming of VLDL/IDL lipid composition provides a plausible mechanistic link between thermogenic adaptation and reduced cardiovascular risk. In addition, the presence of DHA- and EPA-enriched VLDL particles may increase the systemic availability of substrates for the generation of specialized pro-resolving lipid mediators in peripheral tissues, including thermogenic adipose depots and the vascular wall. Through this mechanism, cold-induced remodeling of circulating lipoproteins may not only influence lipid transport and storage but also actively shape inflammatory resolution and tissue-specific adaptive responses.

Importantly, in contrast to obesity, where sustained adipose-to-liver lipid flux culminates in TG accumulation and progressive steatosis, cold exposure is characterized by a remodeling of hepatic lipid composition that preserves the liver from pathological lipid accumulation despite elevated lipid influx. This protection is driven by a biased increase in the biosynthesis of anti-steatotic n-3 HUFAs through desaturase pathways under cold conditions. These findings suggest that endogenous pathways controlling HUFA production are dynamically regulated and may be biased toward n-3 or n-6 substrates, depending on the context. Regulation of this selective desaturase activity may therefore represent a key mechanism governing the hepatic n-3/n-6 balance, with broader implications for metabolic and inflammatory homeostasis.

## Methods

### Animals

Animal procedures were approved by the Institutional Animal Care and Use Committee of the University of São Paulo (CEUA protocol 68/2021). Twelve-week-old male C57BL/6J mice were obtained from the Central Animal Facility of the University of São Paulo (Ribeirão Preto campus). Animals were housed in the experimental animal facility of the Department of Pharmacology (FMRP-USP) under controlled environmental conditions (23 ± 2 °C; 12-h light–dark cycle; relative humidity 45–55%) with ad libitum access to water and standard chow diet (Nuvilab CR-1®, Sogorb Inc.).

Experiments performed at the Joslin Diabetes Center were approved by the Institutional Animal Care and Use Committee at the Joslin Diabetes Center and were conducted using C57BL/6J mice (purchased from Jackson Laboratories), fed with normal chow diet (Mouse Diet 9F, PharmaServ) or HFD (Research Diets, D12492) for 10 weeks. Caloric composition of chow diet consisted of 23% protein, 21.6% fat, and 55.4% carbohydrates, while HFD had 20% protein, 60% fat, and 20% carbohydrates. Mice were watered with either tap water or 30% (wt/v) fructose solution in water, as described by^47^. Fat-1 mice were kindly donated by Dr. Jing X. Kang. After weaning, Fat-1 mice were fed AIN-76A w/10% Corn Oil Diet (5T9W) (Scott Pharma, Inc) until their euthanasia procedure with 12 weeks of age.

For cold-exposure experiments, mice were divided into two groups: control animals maintained at thermoneutrality (30 °C) for 7 days and mice exposed to cold (5 °C) for 7 days. Mice exposed to cold were housed individually in a temperature-controlled chamber with free access to food and water. Animals were euthanized under fed conditions at the end of the acclimation period, with thermoneutral and cold-exposed animals processed in paired fashion on the same day. Euthanasia was performed as described above. Plasma, liver, perigonadal white adipose tissue (pgWAT), subcutaneous white adipose tissue (scWAT), and brown adipose tissue (BAT) were collected for gas chromatography, lipidomics, and other bioenergetic and molecular experiments, as described above.

### Time Course of Cold Exposure in Male Mice

C57BL/6J mice were randomly assigned to six groups: a control group (named time zero) maintained under thermoneutral conditions (30 °C) for 7 days, and five experimental groups exposed to cold (5 °C) for 12 h, 1, 3, 5, or 7 days. At the end of the exposure period, mice were euthanized in the fed state and livers were collected for lipid extraction. For qPCR, western blot, and bioenergetic analyses, the same experimental design was used, but only the 0-, 1-, 3-, and 7-day time points were included.

### Gas-Chromatography coupled to mass-spectrometer

The lipid fractions from the tissues (liver, BAT, ingWAT and pgWAT) were obtained following the^48^ method, modified to solid materials, and in a cold platform. Briefly, the tissue was embedded on 1 mL of Chloroform: Methanol (1:1 – v/v) and homogenized using a Polytron. After, samples were centrifuged at 11,000 rpm for 10 min at 4 °C, and 1 mL of supernatant rescued and added 1 mL of NaOH in methanol, 30 µL of BHT, 50 µL of internal standard (tridecanoic acid – 99% purchased from Merck®) and homogenized using a vortex. Then, samples were heated in a water bath at 100 °C for 15 minutes and immediately refreshed in water at room temperature. Before, 2 mL of boron trifluoride in a methanolic solution (14%) was added, followed by homogenizing and heating at 100 °C for 20 seconds. After refreshing, was added 1 mL of isooctane and vortex. We added 5 mL of saturated NaCl, resting until phase separation. The supernatant was dried using nitrogen. Samples were resuspended with 150 µL of isooctane and followed to the mass-spectrometer.

The chromatographic running was carried out using a GCMS-QP2010 from Shimadzu (Tokyo, Japan), with a silica column Stabilwax (30 m x 0.25 mm, and 0.25 μm of internal diameter) purchased from Restek®. Ultrapure Helium was adopted as running gas (1.3 mL/min). Using an automatic injector (AOC-20i), 1 mL of samples were injected, in a ratio of 1:10 (split). The chromatographic conditions were in accordance to Cintra et al., (2006)^49^, established as 250 °C of injector temperature; oven beginning at 80 °C following 5 °C/min until 175 °C, and 3 °C/min until 230 °C, maintaining for 20 minutes. The conditions of mass spectrometer was established as ionization voltage 70 eV, ionization source was at 200 °C, maintaining full scanning mode with amplitude between 35-500 m/z, and 0.2 seconds by scanning.

### VLDL fractionation for lipidomics analysis

VLDL and IDL were isolated from 770 µL of plasma previously adjusted with potassium bromide (KBr) to a final density of 1.21 g/mL. For gradient preparation, 575 µL of saline solution with a density of 1.063 g/mL were carefully superimposed on the plasma in ultratransparent tubes (Beckman, Brea, CA, USA), using a peristaltic mini pump to minimize mixing between layers (Thomas Scientific, NJ, USA). Then, 575 µL of saline solution with a density of 1.019 g/mL and 500 µL of saline solution with a density of 1.006 g/mL were successively added. The densities of the solutions were confirmed by densimetry (Easy D40, Mettler Toledo, São Paulo, Brazil). The gradient was ultracentrifuged at 55,000 rpm, 15 °C, for 17 h 30 min, in a TLS-55 rotor, using a Beckman Optima TLX microultracentrifuge (Brea, CA, USA). At the end of centrifugation, the fractions were collected sequentially from the top of the tube, obtaining VLDL + IDL (575 µL), LDL (650 µL) and HDL (700 µL).

### Lipidomic analysis

Total lipids were extracted from liver tissue using a modified methyl tert-butyl ether (MTBE) protocol. Briefly, liver tissue (200 mg) was homogenized in 500 µL of buffer containing 10 mM phosphate-buffered saline (PBS) and 20 μM deferoxamine (pH 7.0) using a Polytron electric homogenizer. An aliquot of the homogenate (250 µL) was transferred to a 2 mL tube and mixed with 550 µL MTBE and 600 µL methanol, followed by vortexing for 60 s. Subsequently, 100 µL of the resulting homogenate was transferred to a new tube, and internal standards (mix 1 and mix 2, 50 µL each) were added together with 300 µL of cold methanol containing 10 μM butylated hydroxytoluene (BHT). After vortexing (30 s), 1 mL MTBE was added and samples were incubated for 1 h at 20 °C in a ThermoMixer with constant agitation (1,000 rpm). Phase separation was induced by addition of 300 µL Milli-Q water, vortexing (30 s), and incubation on ice for 10 min, followed by centrifugation at 10,000g for 10 min at 4 °C. The upper organic phase containing the total lipid extract (∼700 µL) was transferred to a new vial and evaporated under a stream of nitrogen. Dried extracts were reconstituted in 100 µL isopropanol. A 10 µL aliquot from each sample was pooled to generate a quality control (QC) sample as previously described^50^.

Lipidomic analyses were performed using electrospray ionization quadrupole time-of-flight mass spectrometry (ESI–QTOF-MS; TripleTOF® 6600, Sciex) coupled to an ultra-high-performance liquid chromatography system (Nexera UHPLC, Shimadzu). Lipids were separated on a CORTECS® UPLC® C18 column (1.6 μm, 2.1 × 100 mm) maintained at 35 °C with a flow rate of 0.2 mL min⁻¹. For reversed-phase chromatography, mobile phase A consisted of water/acetonitrile (60:40, v/v) and mobile phase B consisted of isopropanol/acetonitrile/water (88:10:2, v/v/v). Mass spectrometry data were acquired in both positive and negative electrospray ionization modes over an m/z range of 200–2000^51^. Data processing was performed using Analyst® v1.7.1 (Sciex) and MetaboAnalyst 5.0.

### Isolation of mature adipocytes and stromal vascular (SVF) fraction from adipose tissues

Perigonadal white adipose tissue (WAT) was dissected, finely minced, and enzymatically digested in Hanks’ balanced salt solution containing type I collagenase (1.5 mg/mL) and dispase II (2.5 U/mL), as previously described (Shamsi & Tseng, 2020). Tissue digestion was performed for 45 minutes at 37 °C under gentle agitation. Following digestion, samples were centrifuged at 500 × g for 10 minutes at 4 °C to separate mature adipocytes from the stromal vascular fraction (SVF). The floating adipocyte layer was carefully collected, while the pellet was retained as the SVF fraction. The adipocyte fraction was passed through a 200 μm cell strainer, washed with PBS containing 3% BSA, and centrifuged at 30 × g for 5 minutes to remove residual debris and contaminants. The SVF pellet was treated with ammonium–chloride–potassium (ACK) lysing buffer to eliminate red blood cells. The resulting cell suspension was filtered through a 40 μm cell strainer, washed with PBS supplemented with 1.5% BSA, and centrifuged at 500 × g for 7 minutes to obtain a purified SVF fraction^52^

### Culture and Treatment of AML12 Cells

AML12 cells were cultured in a 1:1 mixture of Dulbecco’s Modified Eagle Medium/Ham’s Nutrient Mixture F-12 (DMEM/F-12; Cultilab) supplemented with 10% fetal bovine serum, 10 μg/mL insulin, 5.5 μg/mL transferrin, 5 ng/mL sodium selenite, 40 ng/mL dexamethasone, 1% of penicillin-streptomycin, at 37°C in a humidified atmosphere containing 5% CO₂. Cells were seeded in 24-well plates and cultured for 48 h until reaching confluence. Subsequently, cells were treated with 0, 50, or 100 µM docosahexaenoic acid (DHA; Merck) for 48 h. In a separate set of experiments, cells were treated with 0, 1, or 10 µM norepinephrine for 24 or 48 h. Ethanol was used as the vehicle control. After treatment, cells were washed with phosphate-buffered saline (PBS) and immediately lysed with TRIzol Reagent (Thermo Fisher Scientific) for RNA extraction or with Cell Lysis Buffer (CLB; Cell Signaling Technology) for protein extraction. For each condition, cells from two independent wells were pooled. All samples were stored at −80 °C until further analysis. All experiments were performed using up to six independent biological replicates, each derived from a different cell passage.

### Culture and differentiation of 9W lineage cells into white adipocytes

Cells of the 9W lineage were maintained in Dulbecco’s Modified Eagle Medium (DMEM) with high glucose, supplemented with 10% fetal bovine serum (FBS) and 1% penicillin–streptomycin. Cells were cultured at 37 °C in a humidified atmosphere containing 5% CO₂ and maintained under exponential growth conditions until initiation of the differentiation protocol. To induce adipogenic differentiation, cells were seeded at a density sufficient to reach full confluence approximately 48 h after plating. Differentiation was initiated at confluence (day 0), with cells initially maintained in complete culture medium. On day 2, differentiation into white adipocytes was induced by replacing the culture medium with induction medium containing insulin (20 nM), dexamethasone (1 µM), 3-isobutyl-1-methylxanthine (IBMX, 500 µM), and rosiglitazone (1 µg ml⁻¹). Cells were maintained in this medium for 48 h. On day 4, the medium was replaced with maintenance medium containing insulin (20 nM), dexamethasone (1 µM), and rosiglitazone (1 µg mL⁻¹), without IBMX. On day 6, cells were maintained in medium containing insulin (20 nM) and dexamethasone (1 µM). By day 8, cells displayed morphological features consistent with differentiated white adipocytes and were considered ready for subsequent experimental procedures.

### Quantitative real-time PCR (RT–qPCR)

Total RNA was extracted from liver tissue using TRIzol Reagent (Thermo Fisher Scientific, Waltham, MA, USA) according to the manufacturer’s instructions. RNA concentration and purity were determined by spectrophotometry (NanoDrop 2000, Thermo Fisher Scientific). One microgram of total RNA was reverse-transcribed using the High-Capacity cDNA Reverse Transcription Kit (Applied Biosystems, Foster City, CA, USA) in a final volume of 20 μL, following the manufacturer’s protocol. Quantitative PCR was performed using SYBR Green Master Mix (Applied Biosystems) on a QuantStudio 5 Real-Time PCR System (Applied Biosystems). Reactions were carried out in 96-well plates with a final volume of 10 μL containing 5 μL of SYBR Green Master Mix, 0.5 μM of each primer, and 2 μL of diluted cDNA. The thermal cycling conditions were: 95 °C for 10 min, followed by 40 cycles of 95 °C for 15 s and 60 °C for 1 min. Melt curve analysis was performed to confirm amplification specificity.

Relative gene expression was calculated using the 2^−ΔΔCt^ method, with Gapdh and Tbp as the endogenous control. A complete list of primers is provided in Supplementary Table 1.

### Western blot analysis

Liver tissues were homogenized in RIPA buffer supplemented with protease and phosphatase inhibitors. Protein concentration was determined using the BCA assay. Equal amounts of protein (20–40 μg) were resolved by SDS–PAGE and transferred to PVDF membranes. Membranes were blocked in 5% non-fat milk in TBS-T for 1h at room temperature and incubated overnight at 4°C with primary antibodies diluted in blocking buffer. The following primary antibodies were used: anti-SCD1 (rabbit monoclonal, 1:1000, Cell Signaling Technology, Cat# 38672), anti-ChREBP (rabbit monoclonal, 1:1000, Cell Signaling Technology, Cat# 58069), anti-ATGL (rabbit polyclonal, 1:1000, Cell Signaling Technology, Cat# 2138), anti-HSL (rabbit polyclonal, 1:1000, Cell Signaling Technology, Cat# 4107), anti-glycerol kinase (rabbit monoclonal, 1:1000, Abcam, Cat# ab126599), anti-PEPCK-C (mouse monoclonal, 1:1000, Santa Cruz Biotechnology, Cat# G-9), and anti-HSP90 (rabbit monoclonal, 1:1000, Cell Signaling Technology, Cat# 4874). Membranes were incubated with HRP-conjugated secondary antibodies for 1h at room temperature and developed using enhanced chemiluminescence. Band intensities were quantified using ImageJ software (NIH). Protein expression levels were normalized to HSP90 as the endogenous loading control. A complete list of antibodies is provided in Supplementary Table 2.

### Fads2 quantification by ELISA kit

To investigate FADS2 protein expression, enzyme-linked immunosorbent assay (ELISA) was performed using liver, perigonadal white adipose tissue (pgWAT), and subcutaneous white adipose tissue (scWAT) samples. Proteins were extracted from tissue homogenates (15–30 mg) in RIPA buffer (Cell Signaling Technology) containing a protease and phosphatase inhibitor cocktail (Thermo Fisher Scientific).

Samples were then centrifuged at 12,000 rpm for 20 min at 4 °C, and the supernatant was collected. In cases where lipid accumulation was observed in the supernatant, the centrifugation step was repeated. Protein concentration in the supernatants was determined using the Pierce BCA Protein Assay Kit (Thermo Fisher Scientific). Protein samples were subsequently analyzed using the MyBioSource Fish Fatty Acid Desaturase 2 ELISA Kit (MyBioSource, Cat# [MBS766186](https://www.mybiosource.com/mouse-elisa-kits/fads2-fatty-acid-desaturase-2/766186)), following manufacturers’instructions.

### High-resolution respirometry

Fresh liver biopsies (∼5 mg) were used for high-resolution respirometry using an Oxygraph-2k system (O2k, Oroboros Instruments, Innsbruck, Austria). Tissue samples were permeabilized in biopsy preservation solution (BIOPS) (2.77 mM Ca²⁺-EGTA, 7.23 mM EGTA, 6.26 mM MgCl₂·6H₂O, 20 mM taurine, 15 mM phosphocreatine, 20 mM imidazole, 0.5 mM DTT, 50 mM MES potassium salt, and 5.77 mM ATP) containing 0.02% saponin for 5 min on ice. Samples were then transferred to the respirometer chambers containing 2.1 mL of air saturated MiR05 respiration buffer (20 mM HEPES, 10 mM K₂HPO₄, 110 mM sucrose, 20 mM taurine, 60 mM potassium-lactobionate, 0.5 mM EGTA, 3 mM MgCl₂, and 1 g.L⁻¹ fatty-acid-free BSA, pH 7.1) at 37°C. Oxygen consumption rates (OCR) were recorded under different respiratory states. The LEAK (non-phosphorylating) respiration state was measured in the presence of succinate (5 mM), malate (10 mM), and glutamate (10 mM). OXPHOS capacity was assessed following stimulation with ADP (4 mM). The maximum electron transfer system (ETS) capacity was determined after additions of the uncoupler CCCP (1 μM, tritation). Residual oxygen consumption (ROX) was determined following addition of NaCN (1 mM) and subtracted from OCR values. Data analysis was performed using the software DatLab 7.4 (Oroboros Instruments).

### β-oxidation assay

Mitochondrial fatty-acid oxidation capacity was assessed using fresh liver biopsies (∼5 mg). Tissues were permeabilized for 5 min in ice-cold BIOPS containing 0.02% saponin and subsequently transferred to the Oxygraph-2k chambers containing 2.1 mL of MiR05 respiration buffer at 37°C. Oxygen consumption was measured in the presence of malate (10 mM) and ADP (4 mM) and fatty-acid oxidation–dependent respiration was then stimulated by addition of palmitoyl-carnitine (5 mM). Residual oxygen consumption (ROX) was determined following addition of NaCN (1 mM) and subtracted from OCR values. β-oxidation–dependent respiration was calculated by subtracting OCR measured in the presence of malate and ADP from OCR measured after palmitoyl-carnitine addition. Values were normalized to citrate synthase activity.

### Triglycerides quantification in mouse liver

Total hepatic lipids were extracted using a NP-40–based method. Liver tissues were weighed and homogenized in 500 μl of an aqueous solution containing 5% (v/v) NP-40. Homogenates were heated at 100 °C until the solution became turbid (∼5 min), followed by cooling to room temperature. Samples were centrifuged at 14,000 rpm for 2 min, and the supernatant containing the lipid fraction was transferred to a new tube. Lipid quantification was performed using a commercial enzymatic kit (Labtest). Briefly, 2 μl of the extracted supernatant was mixed with 200 μl of reagent 1 from the kit in a microplate. Blank wells containing only reaction reagent were included to correct for background signal. Lipid concentrations were determined according to the manufacturer’s instructions.

### Citrate synthase activity

Citrate synthase (CS) activity was measured in samples recovered from the Oxygraph chambers following respirometry assays. The chamber contents were collected and centrifuged at 5,000g for 10 min. The supernatant was discarded, and the resulting pellet was washed three times with 1× HBSS buffer (0.4 mg ml⁻¹ KCl, 0.06 mg ml⁻¹ KH₂PO₄, 0.35 mg ml⁻¹ NaHCO₃, 8 mg ml⁻¹ NaCl, 0.09 mg ml⁻¹ Na₂HPO₄·7H₂O, and 1 mg ml⁻¹ glucose, pH 7.2) to remove residual bovine serum albumin. Pellets were then homogenized in lysis buffer containing 150 mM NaCl, 10 mM Tris-HCl, 1 mM EDTA, and 1% Triton X-100 supplemented with a commercially available protease inhibitor cocktail. Homogenates were centrifuged at 10,000g for 15 min at 4 °C, and the supernatant was collected for protein quantification using the Bradford assay. For the CS activity assay, 5 μg of total protein was added to 200 μl of reaction buffer containing 50 mM Tris-HCl (pH 8.0), 100 μM 5,5′-dithiobis (2-nitrobenzoic acid) (DTNB), and 62.5 μM acetyl-CoA. The reaction was initiated by addition of oxaloacetate (312.5 μM) and monitored for 3 min at 412 nm using a microplate reader. Citrate synthase activity was determined from the slope of the absorbance increase over time.

### Statistical analysis

Statistical analyses for group comparisons were performed using GraphPad Prism (v.9). Data are presented as mean ± s.e.m., unless otherwise indicated. The value of n represents independent biological replicates (individual animals or independently prepared samples), as specified in the corresponding figure legends.

Comparisons between two groups were performed using a two-tailed unpaired Student’s t-test. For experiments involving more than two groups, statistical significance was assessed using one-way ANOVA followed by false discovery rate (FDR) control using the two-stage linear step-up procedure of Benjamini, Krieger and Yekutieli. A P value < 0.05 was considered statistically significant.

Lipidomic and multivariate analysis were performed in R. Lipidomic data from either GC/MS or LC-MS/MS experiments was analyzed using linear models via the limma package^53^. To account for the experimental design, a model matrix was generated based on tissue type and thermal condition. Intra-group correlation was estimated using duplicateCorrelation^54^ and incorporated into the model fit to account for sample dependencies. Specific cold-induced changes within each tissue (BAT, iWAT, pWAT, and liver) were extracted using predefined contrasts. Significance was stabilized via Empirical Bayes moderation^54^, and p-values were adjusted for multiple testing using the Benjamini–Hochberg (FDR) procedure. An adjusted P-value ≤ 0.05 was used as the threshold for significance. Multivariate structure was explored using randomized principal component analysis via rsvd^55^. Data visualization was performed in R using ggplot2^56^and ComplexHeatmap^57^.

## Data Availability

The data that support the findings of this study are available from the corresponding author upon reasonable request. Source data underlying the figures are provided with this paper.

## Acknowledgements

We thank the members of our laboratories for helpful discussions and technical assistance throughout this study. We also acknowledge the support of the core facilities at the University of São Paulo, Ribeirão Preto campus, and at the Joslin Diabetes Center, for their assistance with experimental procedures and instrumentation. This work was supported by grants from Sao Paulo Research Foundation (FAPESP): 2017/08264-8, 2019/26008-4 and 2023/13695-9 (L.O.L). 2021/01311-6 (V.A.V.). 2023/17809-9 (G.S.C.). 2024/10752-4 (G.G.S.). 2018/25053-3 (R.C.G.). 2024/20995-1, 2024/07895-8, and 2021/08354-2 (M.A.M.).

## Author contributions

Conceptualization: L.O.L., V.A.V., G.S.G., G.S.C., R.C.G.; Methodology standardization and experiments: V.A.V., G.S.G., G.S.C., R.C.G., C.S., R.G.C., M.R.S., C.O.R., J.M.A, T.T.G., I.B., C.F.B.L., S.S., M.Y.Y.; Resources: L.O.L., Y.H.T., M.A.M., C.R.K, S.S., D.E.C., A.C.S., J.X.K.; Data analysis and bioinformatics: T.M., M.H.A.C., L.G.G. M.Y.Y.; Writing - original draft: L.O.L., V.A.V., G.S.G., G.S.C., R.C.G.; Writing - review and editing: Y.H.T., M.A.M., C.R.K, S.S., D.E.C., A.C.S., J.X.K., T.M., M.Y.Y.; Project administration: L.O.L.; Funding acquisition: L.O.L., Y.H.T., M.A.M..

