## Supplementary Figures and Tables for "Cold acclimation reprograms hepatic lipid composition toward n-3 HUFAs to uncouple adipose-derived lipid flux from steatosis"

A pgWAT

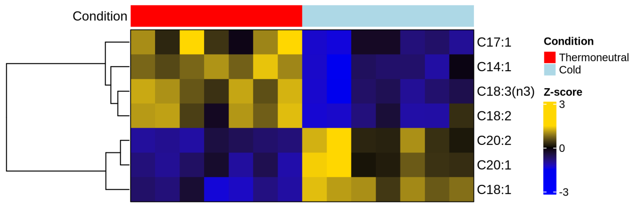


B ingWAT

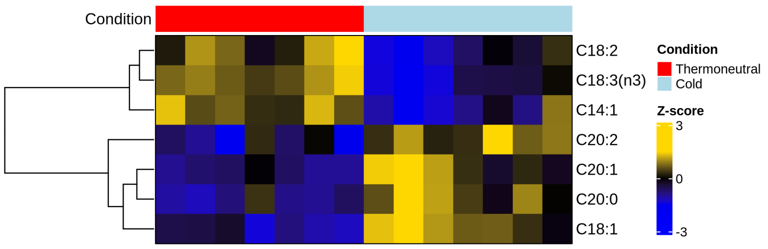


C BAT

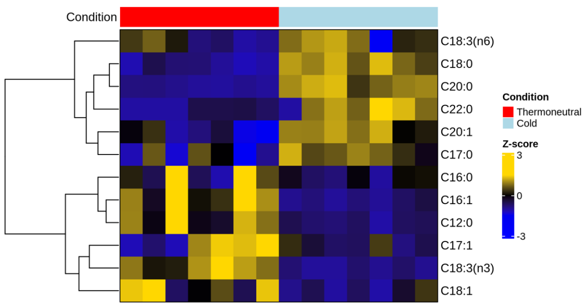


**Supplementary Figure S1 | Tissue-specific fatty acid profiling across adipose depots following cold exposure.**
**A-C,** Heat maps showing the relative quantification of the significantly altered fatty acids quantified by gas chromatography in **(a)** perigonadal white adipose tissue (pgWAT), **(b)** inguinal white adipose tissue (ingWAT), and **(c)** brown adipose tissue (BAT) following 7 days of cold exposure. Each column represents individual biological replicates (n=7).

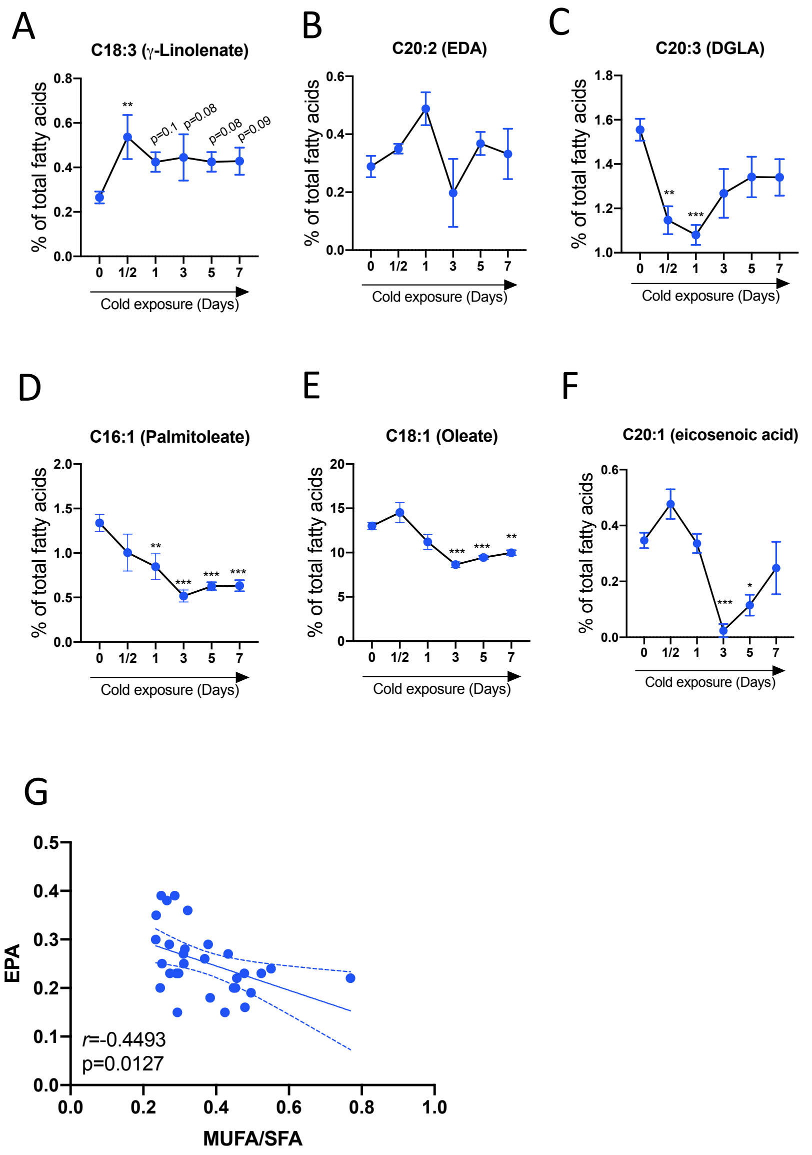


**Supplementary Figure S2 | Cold exposure time course reveals changes in fatty acid composition over the time**

**A-C,** Quantification of n-6 PUFAs γ-linolenate (C18:3, n-6) and eicosadienoic acid (C20:2, n-6) and dihomo-γ-linolenic acid (C20:3, n-6) in the livers collected from mice during the cold exposure time-course.

**D-F,** Quantification of the most abundant MUFAs palmitoleate (C16:1), oleate (C18:1) and eicosenoic acid (C20:1) in the livers collected from mice during the cold exposure time-course. n=4-6 mice per group. Data are expressed as mean ± S.E.M. One-way ANOVA with false discovery rate control using the two-linear stet-up procedure of Benjamini, Krieger and Yekutieli.

**G**, Pearson correlation between the SCD1-dependent desaturation index (MUFA/SFA) and the levels of EPA. The correlation coefficient (*r)* and corresponding two-tailed *P* value were calculated using Pearson's method.

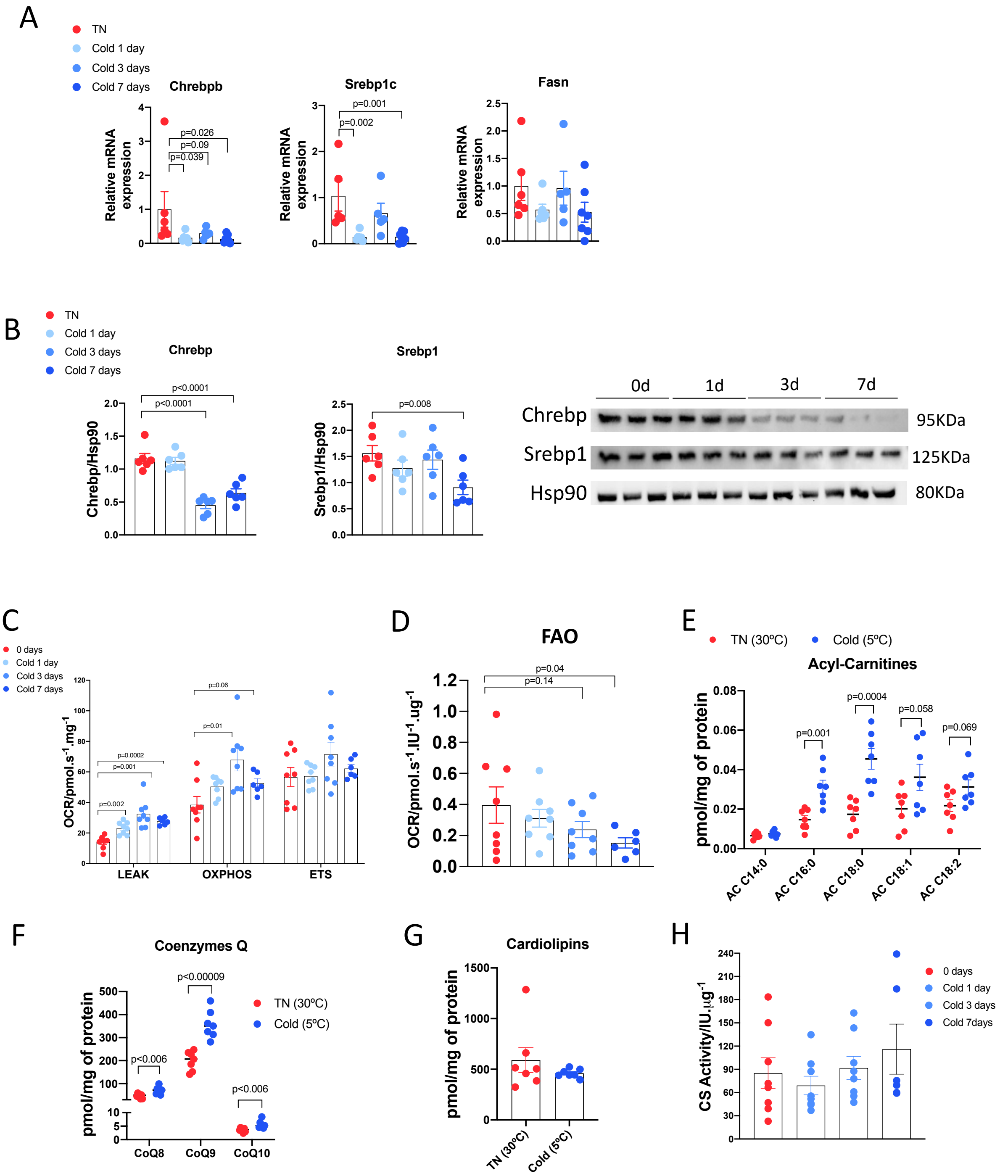


**Supplementary Figure S3 | Cold exposure induce time-dependent reduction in the expression of DNL markers.**

**A,** mRNA expression of DNL markers Chrebpb, Srebp1c and Fasn in the liver of cold exposed mice at different time-points of cold acclimation. n=5-7 per group.

**B,** Bar graphs expressing normalized protein expression of DNL markers Chrebp, Srebp1, accompanied by the representative western blot images. n=6 mice per group. Data are expressed as mean ± S.E.M. One-way ANOVA with false discovery rate control using the two-linear stet-up procedure of Benjamini, Krieger and Yekutieli.

**C,** High-resolution respirometry (Oroboros O2k™) was used to quantify mitochondrial respiratory function in liver biopsies from mice exposed to cold for 0, 1, 3 and 7 days. Proton leak (LEAK), oxidative phosphorylation capacity (OXPHOS) and maximal electron transport system capacity (ETS) were determined.

**D,** Quantification of oxygen consumption through high-resolution respiration to estimate mitochondrial fatty acid oxidation (FAO) in liver biopsies from mice exposed to cold for 0, 1, 3 and 7 days. Palmitoyl-carnitine and malate were used as substrates for FAO capacity assessment. n=6-8 mice per group. Data are expressed as mean ± S.E.M. One-way ANOVA with false discovery rate control using the two-linear stet-up procedure of Benjamini, Krieger and Yekutieli.
**E-G**, Quantification of (**E**) long-chain Acyl carnitines, Coenzyme-Q levels (**F**) and total cardiolipin levels (**G**) through liquid chromatography-tandem mass spectrometry (LC-MS/MS) in the livers from mice exposed to cold or thermoneutrality. N=7 mice per group. Two-tailed unpaired Student *t*-test.
**H**, Citrate synthase activity assay in liver biopsies from mice exposed to cold for 0, 1, 3 and 7 days. n=6-8 mice per group. Data are expressed as mean ± S.E.M.

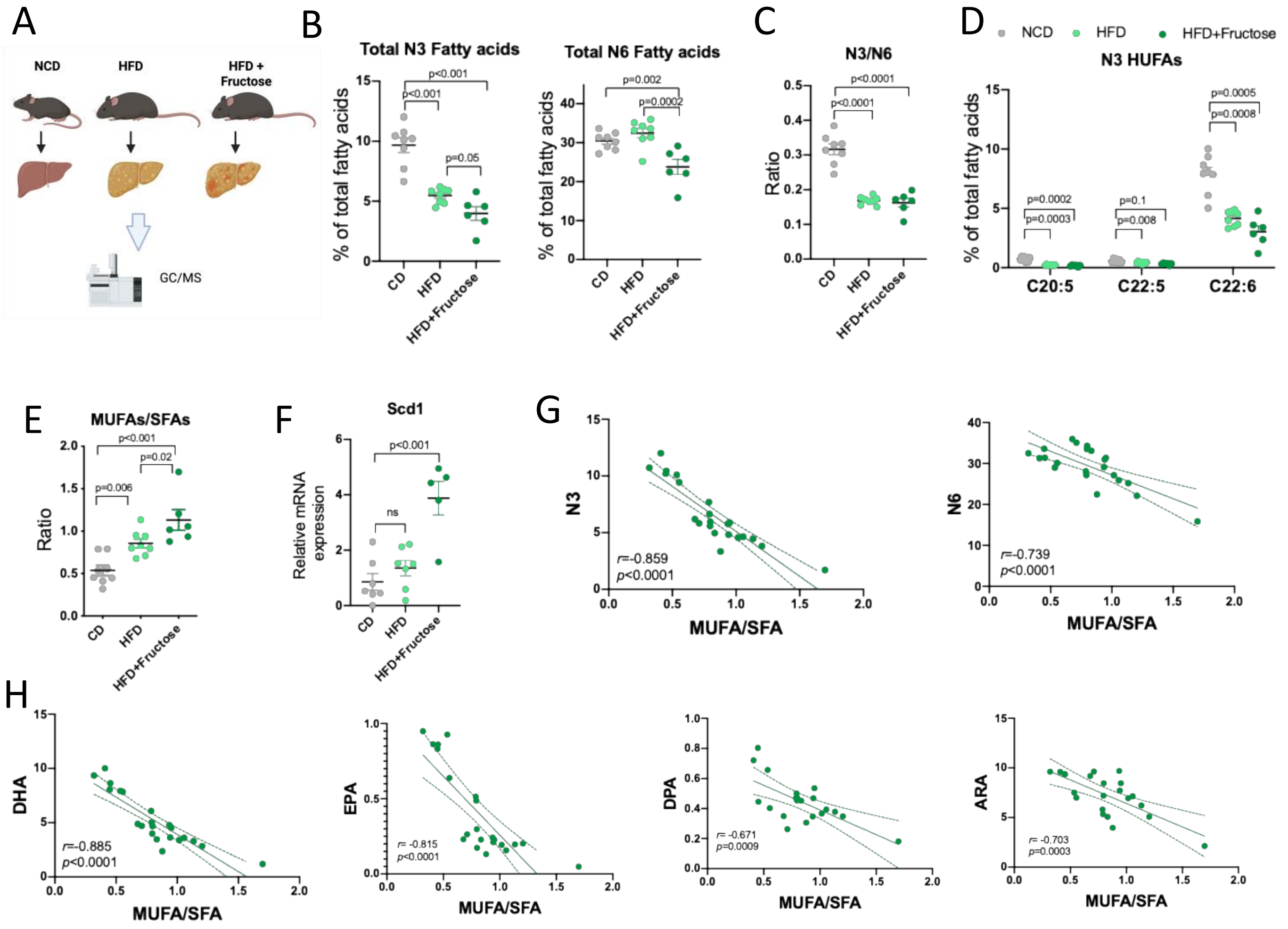


**Supplementary Figure S4 | Inverse correlation between N3-HUFAs and Scd1-desaturation index is maintained in MAFLD models.**

**A,** Schematic representation of the experimental design. Livers from C57BL6J mice, fed with normal chow diet (NCD), high-fat diet (HFD) and HFD + fructose, were harvested and processed for lipid profiling through gas chromatography coupled to mass spectrometry.

**B,** Fatty acid profiling of the sum of n-3 and n-6 fatty acids in the liver of NCD, HFD and HFD + fructose groups.

**C,** Quantification of n-3/n-6 ratio in the liver of the referred groups.

**D,** Quantification of n-3 HUFAs species in the liver of the referred groups.

**E,** Quantification of MUFAs/SFAs ratio in the liver of the referred groups. n=6-8 mice per group.

**F,** mRNA expression of Scd1 in the liver from the referred groups. n=5-7 mice per group. Data are expressed as mean ± S.E.M. One-way ANOVA with false discovery rate control using the two-linear stet-up procedure of Benjamini, Krieger and Yekutieli.

**G,** Pearson correlation between the SCD1-dependent desaturation index (MUFA/SFA) and the levels of total n-3 FAs and n-6 FAs. n= 22.

**H,** Correlation between the MUFA/SFA and the levels of total n-3 FAs (F), n-6 FAs (G), DHA (H), EPA (I), DPA (J) and ALA (K). n= 22. The correlation coefficient (*r)* and corresponding two-tailed *P* value were calculated using Pearson's method.

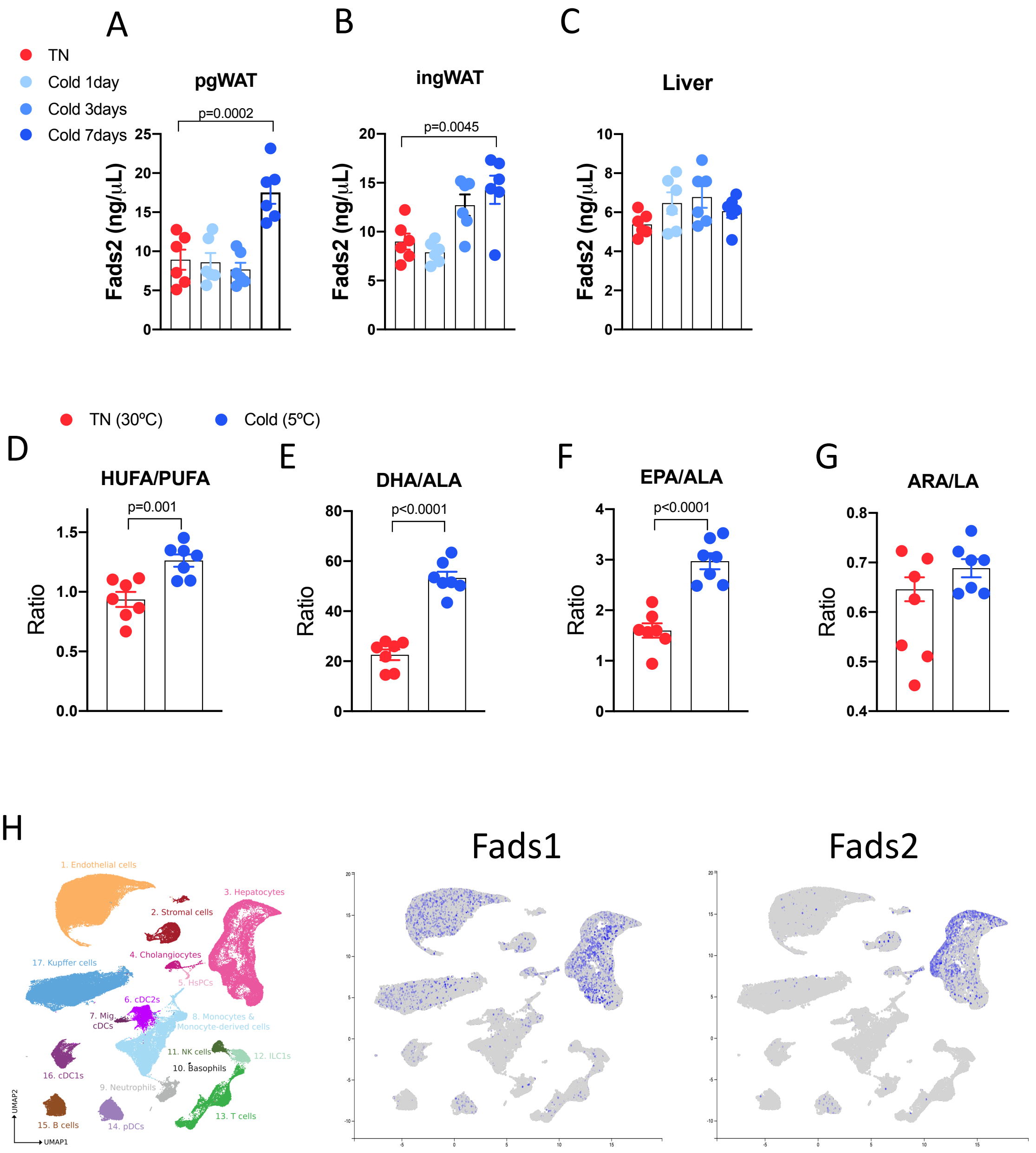


**Supplementary Figure S5 | Fads 1 and 2 expression profile in adipose tissue and liver.**

**A-C,** FADS2 protein (delta-6 desaturase) expression in perigonadal white adipose tissue (pgWAT), inguinal WAT (ingWAT) and liver harvested from mice exposed to cold at time zero, and 1, 3 and 7 days of cold. n=6 mice per group. One-way ANOVA with false discovery rate control using the two-linear stet-up procedure of Benjamini, Krieger and Yekutieli.

**D-G,** Estimative of Fads activity through the quantification of the ratios between: HUFAs/PUFAs (D), DHA/ALA (E) and EPA/ALA (F), in the liver from mice exposed to thermoneutrality or cold. n=7 mice per group. Two-tailed unpaired Student *t*-test.

**H,** Single-cell RNA sequencing data from the publicly available Liver Cell Atlas (https://www.livercellatlas.org). UMAP representation of annotated murine liver cell clusters, with feature plots showing the distribution of *Fads1* and *Fads2* expression across distinct cell populations.

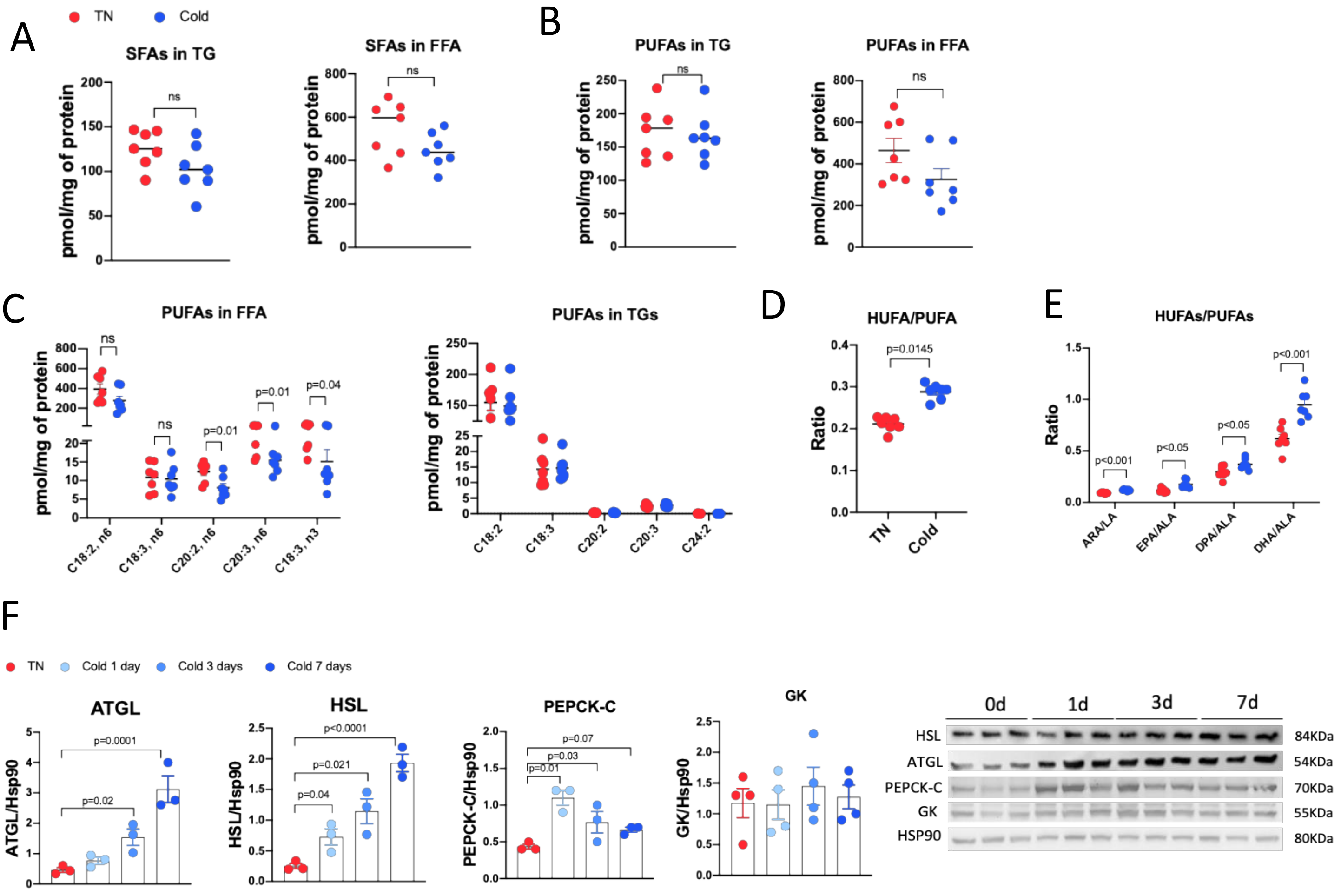


**Supplementary Figure S6 | Fads 1 and 2 expression profile in adipose tissue and liver.**

**A,** Quantification of the sum of saturated fatty acids (SFA) in the FFA and in TG forms in the liver of mice exposed to 5°C or 30°C, through lipidomics assay.

**B,** Quantification of the sum of PUFAs in the form of FFA or TG in the liver of mice exposed to 5°C or 30°C.

**C,** Quantification of individual species of PUFAs in the form of FFA or TG in the liver of mice exposed to 5°C or 30°C.

**D,** Quantification of HUFA/PUFA ratio as FFAs in the liver of mice exposed to 5°C or 30°C.

**E,** Quantification of the ratio between individual HUFAs species and their respective PUFA precursor: ARA/LA, EPA, ALA, DPA/ALA and DHA/ALA. Data are expressed as mean ± S.E.M. n= 7 mice. Two-tailed unpaired Students *t*-test.

**F,** Bar graphs expressing normalized protein expression of triglyceride turnover markers ATGL, HSL, PEPCK-C and GK, accompanied by their respectives representative western blot images. n=3-4 mice per group. One-way ANOVA with false discovery rate control using the two-linear stet-up procedure of Benjamini, Krieger and Yekutieli.

A

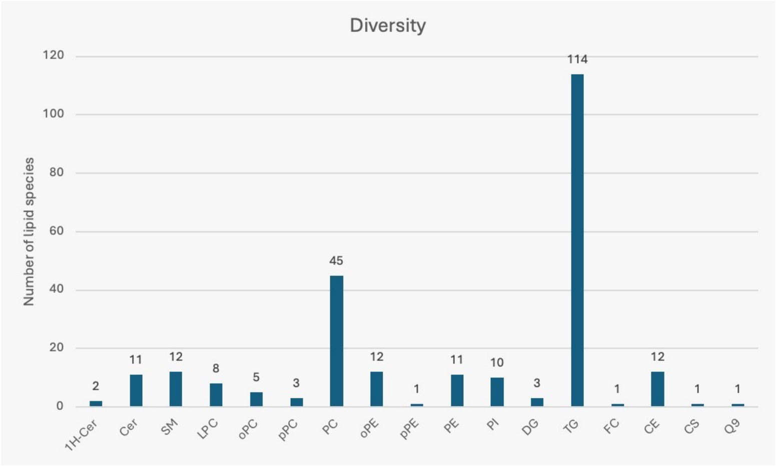


B

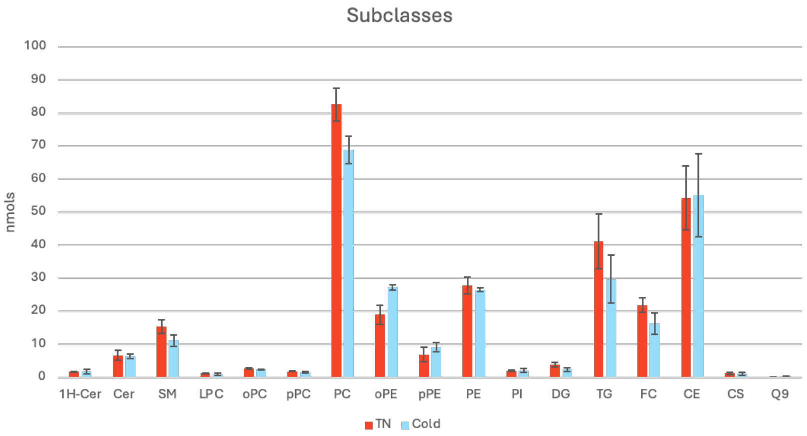


**Supplementary Figure S7 | Lipidome profile in circulating VLDL/IDL fraction**
**A,** Bar graph expressing the diversity of lipid species in the isolated VLDL/IDL compartment. Expressed as the number of lipid species incorporated into each lipid class. N=6.
**B,** Concentration in nmols of each lipid class in the isolated VLDL/IDL compartment. Comparison between cold exposed and thermoneutrality (TN) groups. Data are expressed as mean ± S.E.M. n= 3 mice. Two-tailed unpaired Students *t*-test.

### Supplementary Tables

| Gene | Species | NCBI Reference Sequence (mRNA) | Direction | Sequence (5'→3') |
|---|---|---|---|---|
| **Fads2** | *Mus musculus* | NM_010191.3 (Fads2) | Forward | GAGAAGATGCTACGGATGCCT |
| | | | Reverse | TCAGCAGTCTTCTTCAGGC |
| **Fads1** | *Mus musculus* | NM_007998.4 (Fads1) | Forward | TGAACCCACCAAGAATAAAGCG |
| | | | Reverse | AAGAAGAGGTGGTTGGCCTT |
| **Gapdh** | *Mus musculus* | NM_008084.3 | Forward | AGGTCGGTGTGAACGGATTTG |
| | | | Reverse | TGTAGACCATGTAGTTGAGGTCA |
| **Chrebpb** | *Mus musculus* | NM_001287103.2 | Forward | TCTGCAGATTCGCG |
| | | | Reverse | CTTGTCCAGCATAGCAAC |
| **Srebp1c** | *Mus musculus* | NM_011480.4 (Srebf1) | Forward | TGACCCGGCTATTCCGTGA |
| | | | Reverse | CTGGGCTGAGCAATACAGTTC |
| **Scd1** | *Mus musculus* | NM_009127.4 (Scd1) | Forward | TTCTTGCGATACACTCTGGTGC |
| | | | Reverse | CGGGATTGAATGTTCTTGTCGT |
| **Tbp** | *Mus musculus* | NM_013684.4 (Tbp) | Forward | ACCCTTCACCAATGACTCCTATG |
| | | | Reverse | TGATGACTGCAGCAAATCGC |

Supplementary Table 1 | Primer sequences used for quantitative real-time PCR

| Target | Host | Clonality | Company | Catalog number | Dilution | Application |
|---|---|---|---|---|---|---|
| SCD1 | Rabbit | Monoclonal (C12H5) | Cell Signaling Technology | 38672 | 1:1000 | WB |
| ChREBP | Rabbit | Monoclonal | Cell Signaling Technology | 58069 | 1:1000 | WB |
| ATGL | Rabbit | Polyclonal | Cell Signaling Technology | 2138 | 1:1000 | WB |
| HSL | Rabbit | Polyclonal | Cell Signaling Technology | 4107 | 1:1000 | WB |
| Glycerol Kinase | Rabbit | Monoclonal (EPR6567) | Abcam | ab126599 | 1:1000 | WB |
| PEPCK-C | Mouse | Monoclonal (G-9) | Santa Cruz Biotechnology | G-9 | 1:1000 | WB |
| HSP90 | Rabbit | Monoclonal | Cell Signaling Technology | 4874 | 1:1000 | WB |

Supplementary Table 2 | Antibodies used for Western blot analysis
